# Deciphering the Molecular Mechanisms of BPTF Interactions with Nucleosomes via Molecular Simulations

**DOI:** 10.1101/2025.02.25.640153

**Authors:** Ryan Hebert, Jeff Wereszczynski

**Affiliations:** Department of Physics, Illinois Institute of Technology, Chicago, USA; Departments of Physics and Biology, Illinois Institute of Technology, Chicago, USA; Center for Molecular Study of Condensed Soft Matter, Illinois Institute of Technology, Chicago, USA

## Abstract

Many transcription factors regulate DNA accessibility and gene expression by recognizing post-translational modifications on histone tails within nucleosomes. These interactions are often studied *in vitro* using short peptide mimics of histone tails, which may overlook conformational changes that occur in the full nucleosomal context. Here, we employ molecular dynamics simulations to investigate the binding dynamics of the PHD finger and bromodomain of BPTF, both in solution and bound to either a histone H3 peptide or a full nucleosome. Our results show that BPTF adopts distinct conformational states depending on its binding context, with nucleosome engagement inducing compaction of the multidomain structure. PHD finger binding displaces the H3 tail from DNA, increasing H3 tail flexibility while promoting compensatory binding of the H4 tail to nucleosomal DNA. This redistribution of histone-DNA contacts weakens overall hydrogen bonding with DNA, suggesting localized destabilization of the nucleosome core. Despite electrostatic repulsion limiting direct reader-DNA contacts, strong Van der Waals interactions with the H3 tail stabilize binding. Our results provide atomistic insight into how BPTF engagement modulates nucleosome structure and may facilitate chromatin remodeling.

## 1 Introduction

DNA compaction is crucial for packaging eukaryotic DNA within the cell nucleus.^1^ The human genome contains approximatly 2 meters of DNA, which must be densely packed into a nucleus just 5 to 20 µm across.^2,3^ DNA is a highly electronegative polymer, with each base pair carrying a negative charge in the phosphate backbone. Organization of DNA into chromatin *in vivo* thus requires interactions with a large number of cationic ions and proteins to neutralize this charge.^4^ The first step in eukaryotic DNA compaction is the formation of complexes with eight histone proteins.^5,6^ Approximately 147 base pairs of DNA wrap around a disk formed of an (H3/H4)_2_ tetramer and two H2A/H2B dimers to form the nucleosome core particle (NCP).^7^ Connecting each NCP are 10 to 90 base pairs of linker DNA, and groups of connected NCPs are joined into chromatin fibers.^6,8^ The specific arrangement of nucleosomes within the fiber and thus its diameter vary, with zigzag and solenoid arrangements each observed.^9^ On a scale of hundreds of millions of DNA base pairs, the chromatin fibers construct chromosomal territories and chromosomes.^10^

One effect of this multilevel organization of DNA compaction is to modulate the accessibility of the genome for transcription.^11^ The formation of nucleosomes and chromatin decreases transcriptional activity by blocking the initiation of RNA polymerase II,^12,13^ and for eukaryotic organisms, transcription factors are required for a promoter region to become active.^14^ Although many transcription factors compete with histones for access to DNA,^15–17^ some have reader domains that recognize post-translational modifications on histones.^18,19^ This recognition allows them to target promoters or regulatory regions marked by particular chromatin states.

Histones are composed of disordered N-terminus “tails” and bundles of three alpha helices connected by two loops.^20–22^ Strong electrostatic interactions between phosphates, lysines, and arginines create an interface between the helices and turns of the histone cores with DNA, compacting the negatively charged DNA and the positively charged histones.^23,24^ The histone tails contain about 25% of the total histone mass and are also rich in lysine and arginine, leading to a similar strong DNA association on the exterior of the NCP.^25,26^ Along these tails are many sites for PTMs, among the most common being lysine acetylation and lysine methylation.^27^ PTMs on histones are also markers of active transcriptional sites, such that the histone code is hypothesized to be determinative of gene expression along with the genetic code.^18,28^ Part of the mechanism lies in the altered charge of the amino acids to which PTMs are applied, decreasing the strength of the DNA-histone tail association and increasing the likelihood that a reader may bind to the tail.^29^ However, NMR experiments and molecular dynamics (MD) simulations show that histone tails associate preferentially with DNA in the NCP to the exclusion of PTM readers in mononucleosomes.^30,31^ Histone tails are subject to intermolecular forces in chromatin that are absent in mononucleosome systems, which may free them for interactions with reader domains.^32^

One common reader domain motif is the plant homeodomain (PHD) finger, featuring a three sided cage that recognizes PTMs and zinc binding domains.^33,34^ PHD fingers are found in more than 100 human proteins, often accompanied by other chromatin binding domains, and most commonly bind to lysine 4 on histone H3 in various methylation and acetylation states.^35,36^ A bromodomain is another reader with its own acetylated lysine binding pocket and features four parallel alpha helices connected by loop or loop-helix-loop regions.^37,38^ Bromodomain PHD Finger Transcription Factor (BPTF) is an example of a protein containing both domains - a PHD finger recognizing trimethylated lysine 4 on H3 (H3K4me3) and a bromodomain recognizing acetylated lysine 16 on H4 (H4K16ac).^33,39^ H3K4me3 notably marks the beginning of many transcription sites in the genome,^40–42^ making it an important PTM for structural studies of reader domains. BPTF is part of the Nucleosome Remodeling Factor in humans (hNURF), which is involved in neuronal development^43,44^ and catalyzing nucleosome sliding.^45,46^ The other components of hNURF are SNF2L and pRBAP46/48, an ATPase and a histone binding domain, respectively.^43^ Studies have focused on BPTF as a target for drug therapy due to its enhancement of cancerous tissue growth^39,47–49^ in addition to its contribution as part of hNURF in chromatin remodeling.

BPTF reader domains are often studied *in vitro* using short peptides that mimic the histone tail and PTM,^33,50^ which may preclude understanding of the dynamics in the context of full-length nucleosomes and chromatin. Observation of such systems *in vivo* and production of full NCP-reader systems *in vitro* likewise proves difficult.^51^ Here, we have used classical molecular dynamics (MD) simulations to explore the differences in the motions and interactions of a PHD finger and finger-linker-bromodomain fragment of BPTF in solution, with a peptide fragment mimicking H3, and a full-length NCP. MD simulations provide an ideal computational counterpart to *in vitro* experiments, as they can provide atomistic details of nucleosome motions across a wide range of length and timescales in nucleosome systems. Previous simulation studies have shown strong agreement with experimental observations of histone tail behavior, nucleosomal breathing, and structural transitions, supporting the validity of these methods in chromatin contexts.^26,52–56^ Our findings indicate that NCP binding of the BPTF fragment results in different conformation states of the reader protein that are not accounted for when binding to H3-mimicking peptides, as well as a reduction in histone-DNA hydrogen bonding and shift in H3-H4 tail competition when binding to PHD fingers. This suggests a potential mechanism to induce nucleosome sliding shared by PHD finger containing proteins, and that the full dynamics of readers may not be fully captured in studies with short peptides.

## 2 Methods

### 2.1 System construction

NCP models were generated by taking the 1KX5 crystal structure,^7^ remodeling the H3 tails to extend away from the NCP core using UCSF Chimera’s Modeller interface,^57,58^ and introducing a trimethylation to lysine 4 on one of the H3 tails using tleap from the AmberTools suite.^59^ PHD finger and BPTF models were from the 2F6J crystal structure,^33^ using chain A for BPTF and the chain P H3 residues. Using UCSF Chimera’s Modeller interface,^57,58^ residues 1-15 of H3 were remodeled in reader with peptide systems from the six resolved residues in chain P of the 2F6J structure. PHD finger systems were modified from BPTF by removing the bromodomain and loop region after the helical linker beyond residue 61. NCP and reader complexes were created by using the alignment tools in Chimera to align H3 1-6, deleting those residues from the NCP, and splicing the residues from the 2F6J structure in their place. This maintained the H3 binding mode to the PHD finger and the histone core positioning in the NCP. All starting structures are represented in Figure 1.

**Figure 1:**
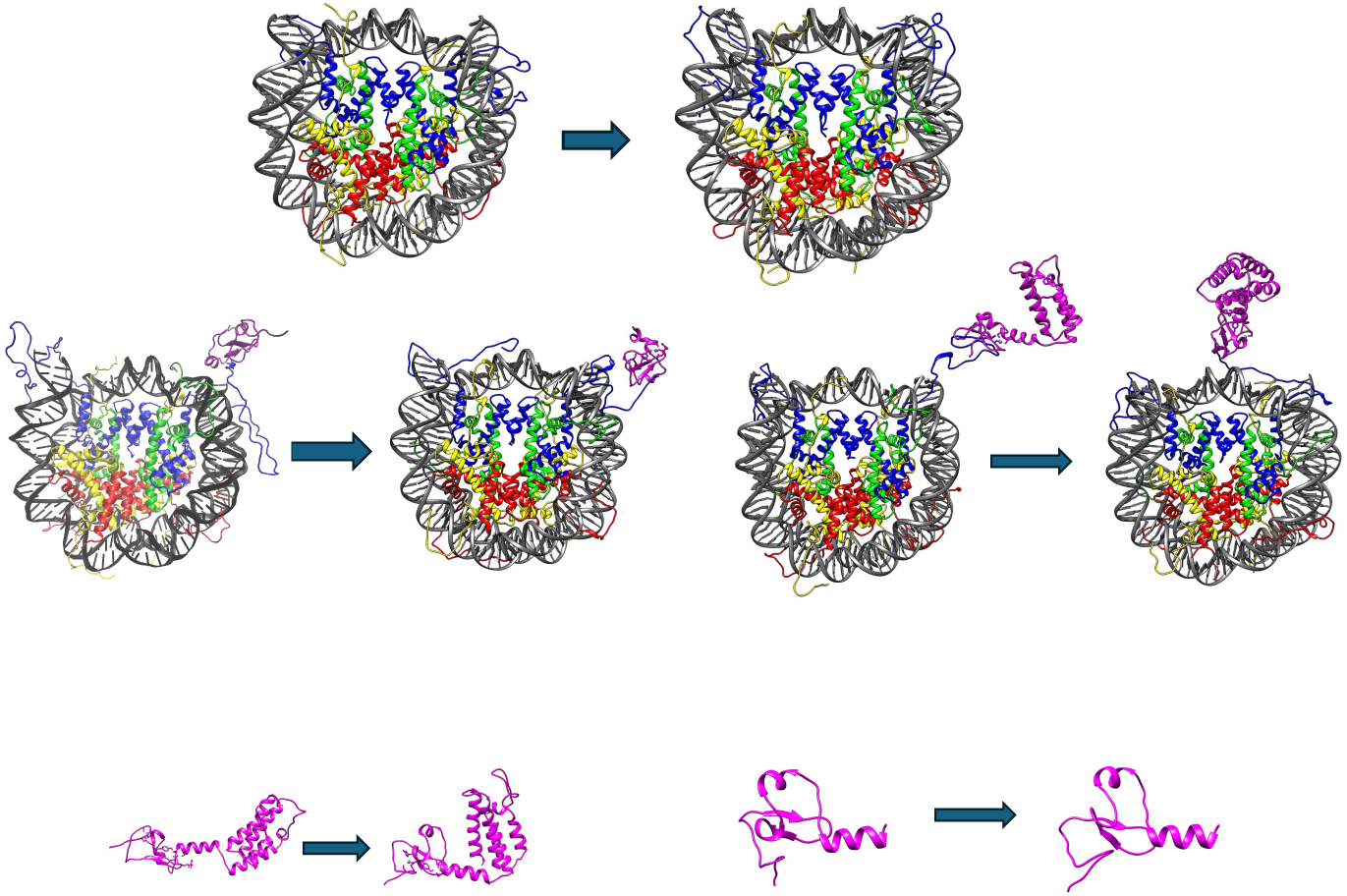
Top left through top right: Sample post-equilibration frames of the simulations for the nucleosome core particle. Middle left through middle right: Post-equilibration frames of the NCP with bound PHD finger and the NCP with bound BPTF fragment. Bottom left through bottom right: Sample post-equilibration frames from simulations of BPTF and the PHD finger. The first in each is the first equilibrated frame from the first simulation and the second is the last frame from the last simulation.

Solvation and generation of parameter and starting coordinate files was performed in tleap and parmed from AmberTools.^59^ All simulations used the ff19SB force field,^60^ an additional library for trimethylated lysine,^61^ OPC water model,^62^ and Li and Merz 12-6 OPC water ions.^63,64^ NCP simulations also used the BSC1 DNA force field.^65^ The BPTF PHD finger contains two zinc metal centers which were paramaterized with the Zinc Amber Force Field.^66^ All but two systems were solvated in water boxes with a 10 Å buffer, neutralized with Na+ ions, and additional Na+ and Cl-ions added to give final ion concentrations of 150 mM NaCl. To prevent self-image interactions, BPTF systems without nucleosomes were solvated in isometric water boxes with the same ion neutralization and final ion concentrations. To decrease computation time, hydrogen mass repartitioning^67^ was employed in parmed, changing the masses of all non-solvent hydrogens to three AMUs and compensating by reducing the mass of the attached heavy atom.

### 2.2 Molecular dynamics simulations

Systems were minimized twice for 10000 steps, swapping from steepest descent to conjugate gradient after 5000 steps. The first minimization applied a 10 kcal/mol · Å*^−^*^2^ harmonic restraint to all heavy atoms and the second minimization had no restraints. Structures were then heated from 5 to 300 K over 5 ps using a Berendsen barostat and Langevin dynamics in an NVT ensemble^68,69^ with 10 kcal/mol · Å*^−^*^2^ harmonic restraints on solute heavy atoms. These restraints were then relaxed by factors of 1/3 every 100 ps in the NPT ensemble and completely removed after 600 ps. All systems after relaxation were simulated in five copies each using pmemd.cuda^59,70–72^ for an additional microsecond in the NPT ensemble at 300 K, 4 fs timesteps, and a 10 Å non-bonded cutoff using local resources and resources provided by the Extreme Science and Engineering Discovery Environment (XSEDE).^73^ Based on root mean square deviation from the first frame (RMSD) data calculated using cpptraj^59^ and observing H3 tail collapse in each trajectory, the first 300 nanoseconds of each simulation were considered equilibration time in analyses. All minimization, heating, relaxation, and simulations were performed using Amber 20.^59^

### 2.3 Simulation Analyses

RMSD, angle measurements between the PHD finger and bromodomain, root mean squared fluctuations (RMSF), hydrogen bonds, and radius of gyration analyses were performed using cpptraj from AmberTools20^59^ on post-equilibration trajectory frames. To measure RMSFs, histones first underwent an RMSD alignment to the heavy atoms in their alpha helices on the first post-equilibration frame. The average atomic positions of the histone heavy atoms was then computed, and RMSFs were then measured referencing these average positions. RMSFs of the PHD finger were measured in a similar manner, aligning to the heavy atoms in the first 59 residues in the PHD finger. Hydrogen bonds were calculated using a donor-acceptor cutoff of 3.0 Å and a cutoff angle of 135°. All intramolecular and DNA base pair hydrogen bonds were excluded.

Solvent accessible surface area was calculated in AmberTools using the linear combinations of pairwise overlaps method^74^ on the H3 tails. Per-residue H3 contacts with DNA and readers, histone and DNA contacts, and measurement of reader atoms within a Debye length of DNA were calculated using MDAnalysis^75,76^ with 4.5 Å cutoffs between heavy atoms for contacts and 7.8 Å cutoffs for the Debye length in 150 mM neutral NaCl.^77^

Contacts across adjacent residues were considered correlated in reporting averages across the H3 tail. Histone and DNA contacts were calculated in a similar manner, where residues 1 - 16 in H2A, 1 - 34 in H2B, 1 - 36 in H3, and 1 - 30 in H4 are considered tails. The remaining residues were considered to be the histone core.

Superhelical locations (SHLs) in a range of ±7 took the first and last 6 base pairs of DNA as locations ±7. Subsequent steps of ±0.5 correspond to the next 5 base pairs, separating the 147 base pair sequence into 29 regions. Entry DNA to the dyad corresponded to SHLs −7.0 to 0.0, and going from the dyad to exit DNA corresponded to SHLs 0.0 to +7.0.

MM/GBSA^78^ results were calculated using MMPBSA.py^79^ in Amber20.^59^ Interactions between readers and NCP used the three trajectory approach with the reader trajectories as the receptor, NCP trajectories as ligand, and reader-NCP trajectories as complexes for inputs. Interactions between H3 and H4 tails with DNA used the single trajectory approach with all but DNA and the interacting tails stripped from the trajectory. All post-equilibration frames of the ligand and receptor trajectories were used in each MM/GBSA analysis, and complex trajectories were run one trajectory at a time with an igb value of 8, corresponding to the mbondi3 radii set.^80,81^ Decorrelation times were determined using the timeseries tool of the Python implementation of the multistate Bennett acceptance ratio (pymbar)^82,83^ on the total energy timeseries of the complex for each calculation.

Angles were measured using vector dot products - one vector connected the centers of mass of α-carbons 7 - 48 in the PHD finger to α-carbons 49 - 56 in the linker, and the second vector connected the centers of mass of α-carbons 137 - 142 and 148 - 153 near the bromodomain binding pocket to centers of mass of α-carbons 125 - 130 and 160 - 165 (see Figure S1).

PHD finger binding pocket volumes were calculated using the sphere method in the Epock plugin for VMD.^84,85^ Trajectories were stripped of all but the PHD finger and aligned to residues 6 - 50 of the H3 peptide bound PHD finger from the 2F6J structure in VMD. 5.0 Å inclusion spheres were centered on the first five H3 peptide residues in the binding pocket.

## 3 Results

### 3.1 Reader Binding Alters H3 Tail Conformation and Dynamics

An array of molecular models was constructed to examine the effects of binding by the BPTF PHD finger and the bromodomain to full-length nucleosomes. These included models of an isolated PHD finger, an isolated PHD-linker-bromodomain complex (referred to as BPTF), a PHD finger and BPTF bound to an H3 peptide, a PHD finger and BPTF bound to a full-length nucleosome, and an isolated nucleosome system (see Figure 1 for examples). Each system was simulated five times for 1.0 µs, giving a total of 35.0 µs of simulation (see Table 1). The H3 tails collapsed in both reader-bound and unbound systems, with both visual inspection and reduced solvent accessible surface area (SASA) showing collapse on a similar timescale of 100–200 ns (Figures S2 and S3).

**Table 1:** A summary of the simulation sampling time of each system.

| System | Simulation Time (ns) | Replicates | Total (ns) |
| --- | --- | --- | --- |
| PHD Finger (PHD) | 1000 | 5 | 5000 |
| PHD + Peptide | 1000 | 5 | 5000 |
| BPTF fragment (BPTF) | 1000 | 5 | 5000 |
| BPTF + Peptide | 1000 | 5 | 5000 |
| H3K4me3 Nucleosome (NCP) | 1000 | 5 | 5000 |
| PHD + NCP | 1000 | 5 | 5000 |
| BPTF + NCP | 1000 | 5 | 5000 |
| Total Simulation Time: |  |  | 35000 |

Reader engagement with the H3 tail remained stable throughout, including during and after tail compaction. Modified H3 tails showed reduced SASA in reader-bound systems, consistent with more extensive interactions with the reader domains (Figure S4). To better understand the nature of this interaction and whether reader binding competes with DNA, we quantified per-residue contacts between the H3 tail, the reader domains, and DNA. Residues H3A1 through H3T6 formed an average of 169 ± 22 and 175 ± 13 contacts with the PHD finger and BPTF, respectively. These values were comparable to those in peptide-bound systems. This suggests that H3 engages the readers in a similar way in the full nucleosome context. Beyond H3T6, reader contacts declined sharply, with fewer than 10 contacts per residue in NCP-bound systems and negligible contacts past H3A7 in systems lacking the full nucleosome (Figure 2). Overall, residues H3A1 through H3A15 formed over 100 additional reader contacts in bound systems. However, this increase in reader engagement was accompanied by a reduction in H3–DNA contacts, highlighting a trade-off between binding to the reader and to DNA.

**Figure 2:**
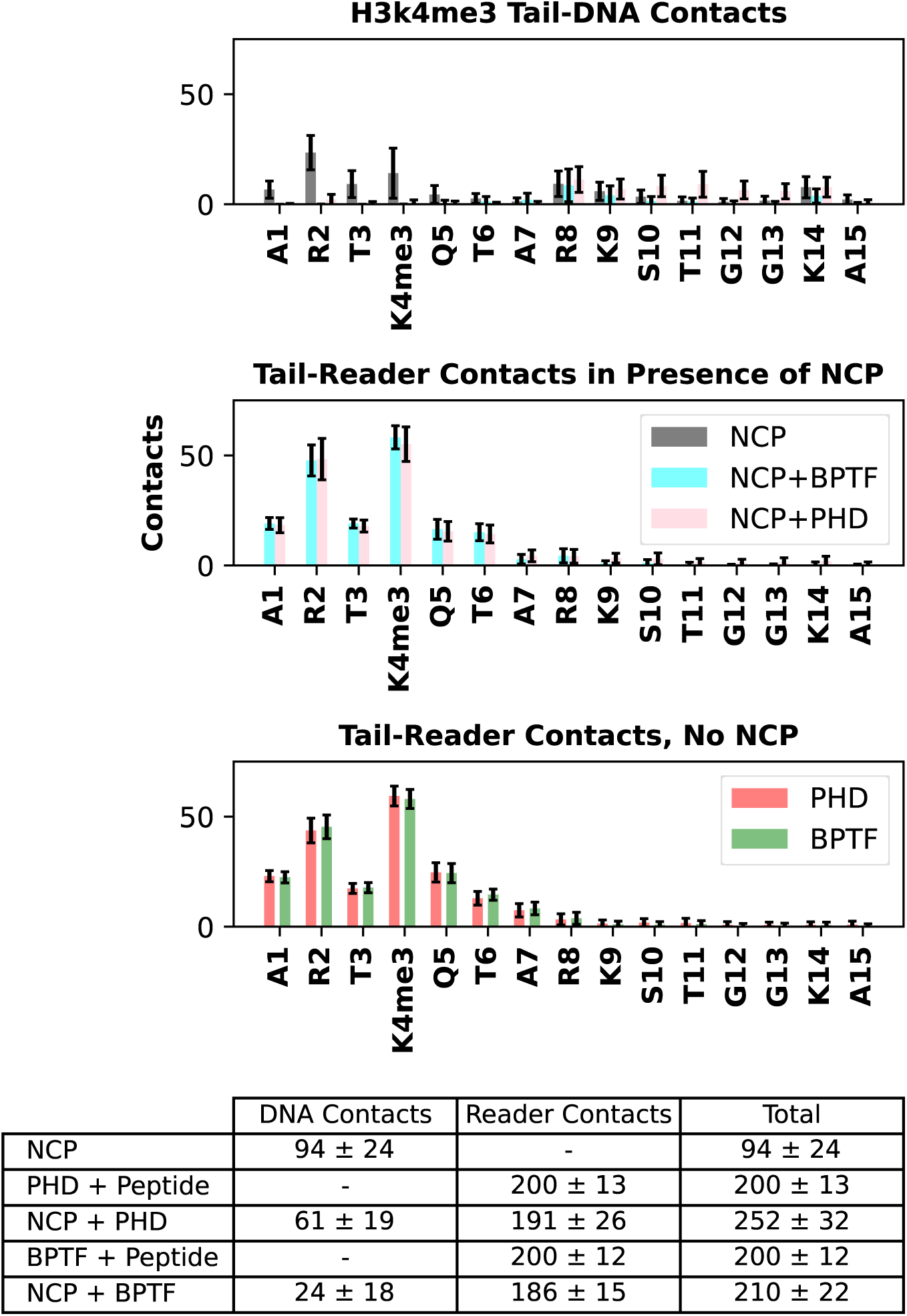
The distribution of contacts over the first 15 residues of the modified H3 tail. Top: Tail contacts with DNA in all NCP systems. Top Middle: Tail contacts with the reader protein in NCP systems. Bottom middle: Tail contacts with the reader protein without the NCP. Bottom: A table summarizing the split in contacts between the reader and DNA. The readers compete with DNA for tail association, with more tail contacts with the reader at the binding pocket and less along the tail. H3 1-15 contacts with the readers shows strong consistency with and without the full NCP.

Reader binding excluded the N-terminal H3 tail residues from interacting with nucleosomal DNA. In unbound NCP systems, residues H3A1 through H3T6 made 60 ± 20 contacts with nucleosomal DNA, but these contacts dropped to near zero in both reader-bound systems (Figure S5). Across the first 15 residues of the H3 tail, total contacts with either the reader or nucleosomal DNA averaged 94 ± 24 in unbound NCPs. In contrast, the same residues formed 252 ± 32 and 210 ± 22 contacts in PHD- and BPTF-bound systems, respectively. These results indicate that the first six residues preferentially associate with the reader. Binding by the larger BPTF complex further reduced H3–DNA engagement, suggesting that reader size amplifies this competitive exclusion.

We evaluated the energetic consequences of reader binding using an MM/GBSA analysis on the H3 tail–DNA interactions. The loss of H3–DNA contacts was accompanied by a decrease in Van der Waals interactions, resulting in weaker binding affinity. In unbound NCPs, the H3K4me3 tail exhibited an average binding energy of –64.6±5.3 kcal/mol. This energy weakened to –45.4±12.6 kcal/mol in PHD-bound systems and to –40.6±6.8 kcal/mol in BPTF-bound systems. Additionally, the presence of the PTM itself appeared to weaken H3’s binding energy by approximately 13 kcal/mol compared to the unmodified H3 tails (Tables S1 & S2).

Per-residue energy contributions showed that the first seven residues in reader-bound systems provided little or no stabilizing interaction with nucleosomal DNA, whereas in unbound tails they contributed between –2.5 and 4 kcal/mol. Beyond this N-terminal region, unbound tails consistently formed more favorable interactions with DNA (Figures S6 & S7). This reduction in stabilizing interactions not only weakened binding but likely enhanced the tail’s conformational freedom.

We then quantified changes in histone tail flexibility by calculating root-mean-square fluctuations (RMSFs) for H3A1–H3A15. Reader binding significantly increased tail mobility, and the larger BPTF complex amplified this effect. In unbound NCPs, RMSFs ranged from 8.5±1.5 to 4.6±0.6 Å, decreasing from the flexible N-terminus toward the histone core. Upon reader binding, maximum RMSFs increased to 11.4±1.5 in the PHD-bound system and 19.4±3.2 in the BPTF-bound system (Figures 3 & S8). These results indicate that reader engagement increases H3 tail motions, and that larger readers cause more pronounced disruption. This combination of increased flexibility and reduced DNA interaction may enhance local accessibility for chromatin regulators and nucleosome-binding proteins.

**Figure 3:**
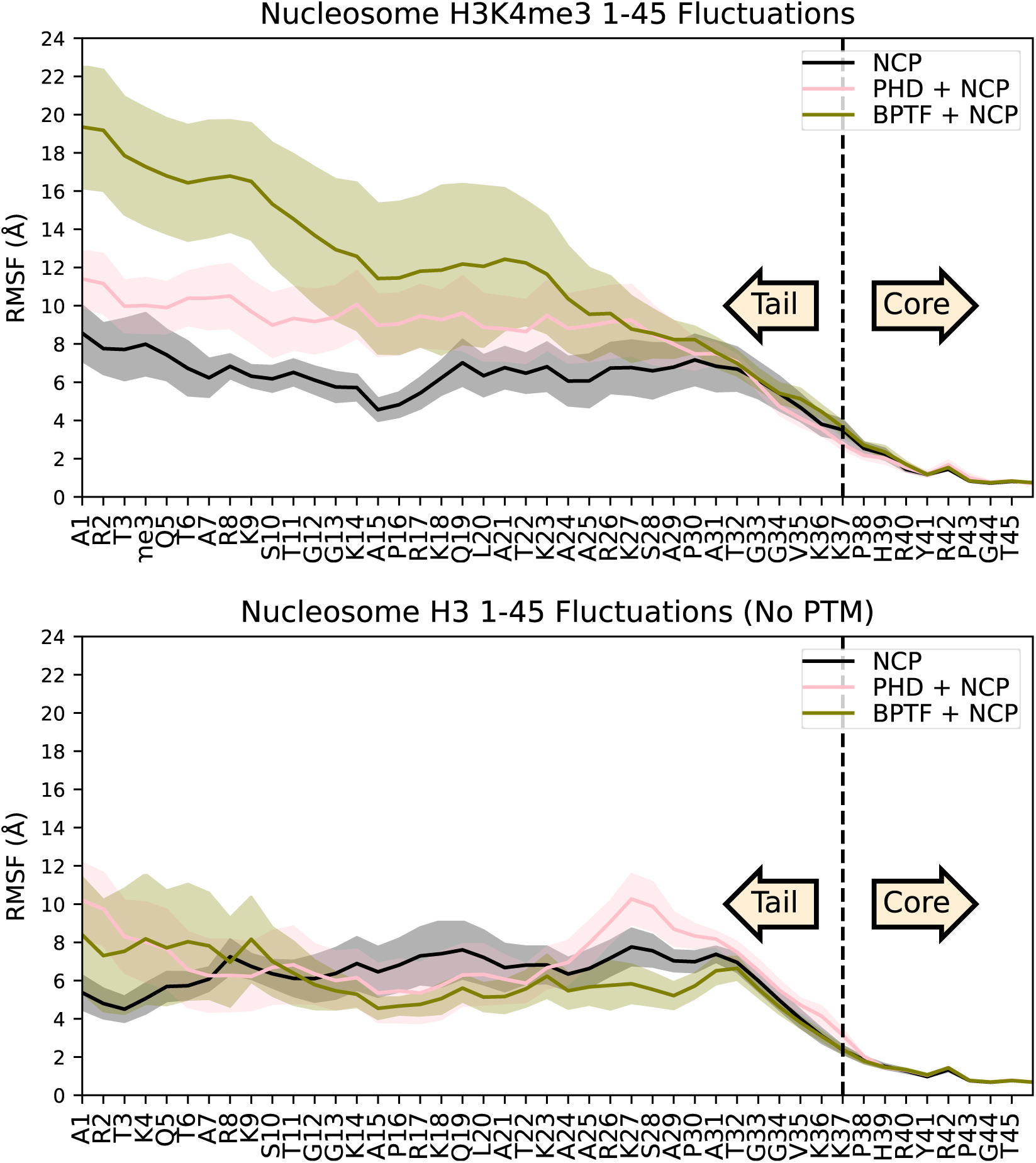
Fluctuations of both H3 histones in each system. In all cases, alignment is to the helices in the H3 core. Top: Fluctuations by residue in the modified H3K4me3 system, where the readers were bound. Trimethylation of lysine 4 and reader domain binding results in an increase in the fluctuations of the H3 tail. The effect is increased by binding the larger reader. Bottom: Fluctuations by residue of the mirror histone H3, where the readers are not bound and there is no PTM.

### 3.2 Binding of H3 Facilitates H4 Interactions with DNA

Having shown that reader engagement displaces the H3 tail from DNA, we next investigated whether this displacement affects H4 tail binding to nucleosomal DNA. We observed substantial overlap in the preferred binding regions of H3 and H4 on the PTM-modified side of the NCP. In contrast, on the unmodified side H3 and H4 tails contacted distinct DNA regions. The overlapping binding sites on the modified side were located near superhelical locations (SHLs) –1.5 to −0.5, corresponding to the base pairs adjacent to the dyad in the direction of the entry DNA (Figures S9-S11). This spatial overlap suggests that the H3 and H4 tails were frequently in proximity and may have competed for DNA access.

In reader-bound systems, H3 tails near the PTM-modified side of the NCP exhibited elevated fluctuations, consistent with displacement from nucleosomal DNA by the reader domains. This displacement exposed regions of DNA that were previously occupied by H3, while H4 tails in the same region showed reduced mobility, suggesting a compensatory gain in H4–DNA interactions (Figures 3 & S12). Similar differences in fluctuations were not observed in the other histones (Figures S13 & S14).

These dynamic changes were accompanied by a redistribution of tail–DNA contacts. On the PTM-modified side, H3 tail contacts with DNA decreased from 206 ± 35 to 149 ± 29 with PHD binding, and further to 134 ± 30 with BPTF binding. In contrast, H4 tails on the same side of the dyad axis increased DNA contacts from 110 ± 26 to 180 ± 31 in the PHD-bound system and to 164 ± 25 in the BPTF-bound systems (Figure 4). This coordinated change indicates a direct mechanistic link: reader binding to H3 reduces its occupancy of the dyad-adjacent DNA, enabling H4 to adopt a more dominant binding role in that same region. Contact levels for H4 on the modified side ultimately approached those on the unmodified side of the NCP, where competition between the tails is minimal.

**Figure 4:**
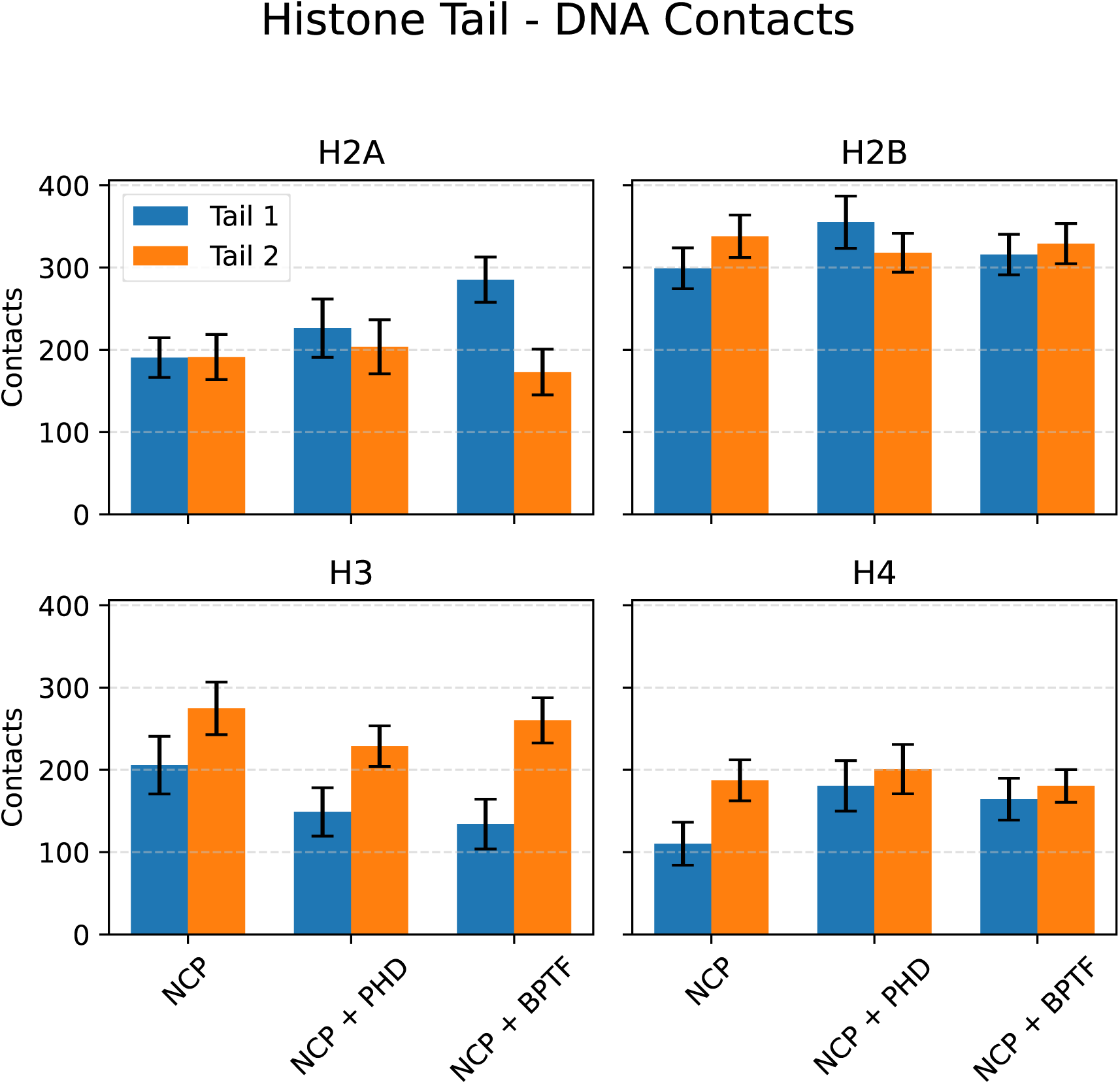
Average atomic contacts between histone tails and DNA. Tail 1 refers to N-terminus histone tails on the reader-bound side of the NCP. Tail 2 refers to N-terminus histone tails on the unbound side of the NCP. Reader binding resulted in a loss of H3 contacts from the bound tail and a corresponding increase in H4 contacts. BPTF binding also resulted in an increase of H2A contacts from tail 1.

Differences in contact frequencies were primarily driven by lysines and arginines, although many residue-level changes fell within the margin of error (Figure S15). Enhanced binding of these positively charged residues to DNA would be expected to reduce H4 tail flexibility, anchoring the tail more tightly to the nucleosomal surface. This restricted mobility is consistent with the reduced RMSFs observed in reader-bound systems (Figure S12). Energetic analyses further supported this mechanism. Near the modified H3 tail, H4–DNA binding energies averaged 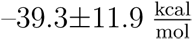 in unbound systems. Upon reader binding to H3, H4–DNA interactions strengthened, reaching 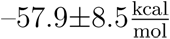 in the PHD-bound system and 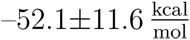 in the BPTF-bound system. These values closely matched H4 binding energies on the unmodified side of the NCP (Tables S3 & S4), mirroring the redistribution of tail contacts and supporting a mechanism in which reader engagement with H3 promotes H4 association with nucleosomal DNA.

These changes in tail positioning and DNA engagement indicate that reader binding not only displaces H3 but also promotes H4 association with nucleosomal DNA. To evaluate the broader consequences of this reorganization, we next examined how reader engagement affects the overall pattern of histone–DNA interactions across the nucleosome.

### 3.3 Reader Binding Shifts Histone-DNA Interactions

Having established that reader binding redistributes H3 and H4 tail contacts, we next quantified how this reorganization affects histone–DNA interactions across the full nucleosome, including H2A and H2B. The overall number of contacts changed little, with only a 2 to 4% difference between unbound and reader-bound systems. However, the distribution of these contacts changed substantially. H3 tails, previously shown to disengage from DNA upon reader binding, lost 103 and 86 contacts in the PHD- and BPTF-bound systems, respectively. These losses were counterbalanced by gains from other histone tails. H4 gained 84 and 48 contacts on average, while H2A increased by 48 and 76 contacts (Figure 5). These findings indicate a cooperative redistribution of histone–DNA contacts, in which loss of H3 binding is compensated by gains from H4 and H2A, maintaining overall nucleosome engagement with DNA.

**Figure 5:**
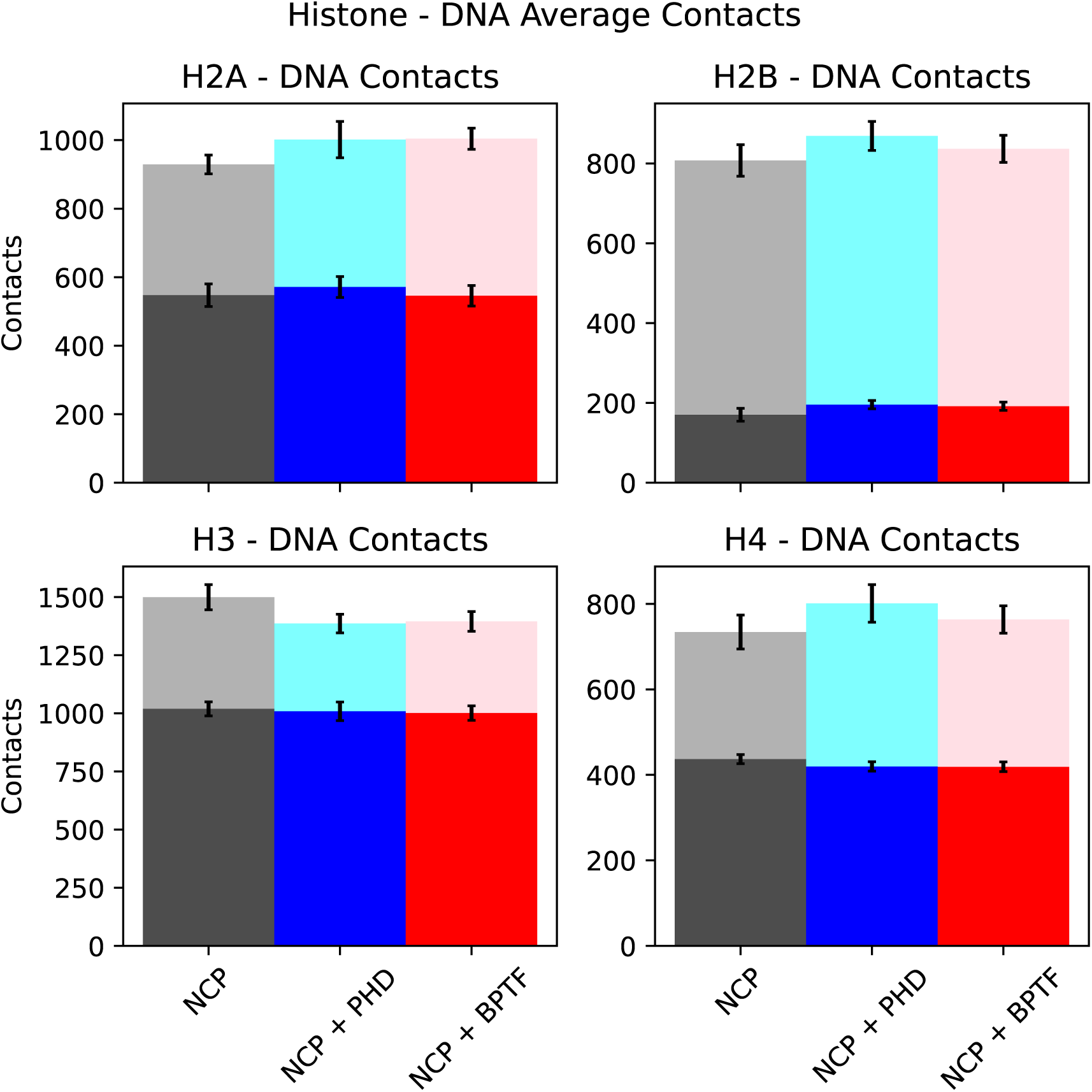
Average atomic contacts between histone tails and histone cores with DNA across the systems with reader domains. Values are stacked with tail contacts on top of core contacts. In both cases histone H3 loses contacts from reader binding, while H2A and H4 gain contacts. Binding BPTF results in a small net loss of H2B-DNA contacts, while PHD binding results in a small net increase. The total number of atomic contacts between histones and DNA in PHD binding systems has a slight net increase, and BPTF binding systems see a net decrease in contacts, indicating a shift in contact distributions.

To determine whether this redistribution also impacted histone-DNA interaction strength, we quantified hydrogen bonding across the histone core. In unbound nucleosomes, H3 histones contributed 49±3 hydrogen bonds to DNA, which fell by 8 in PHD-bound and by 9 in BPTF-bound systems (Figure 6). This decrease occurred in both tail and core regions, suggesting a broad weakening of H3-DNA interactions.

**Figure 6:**
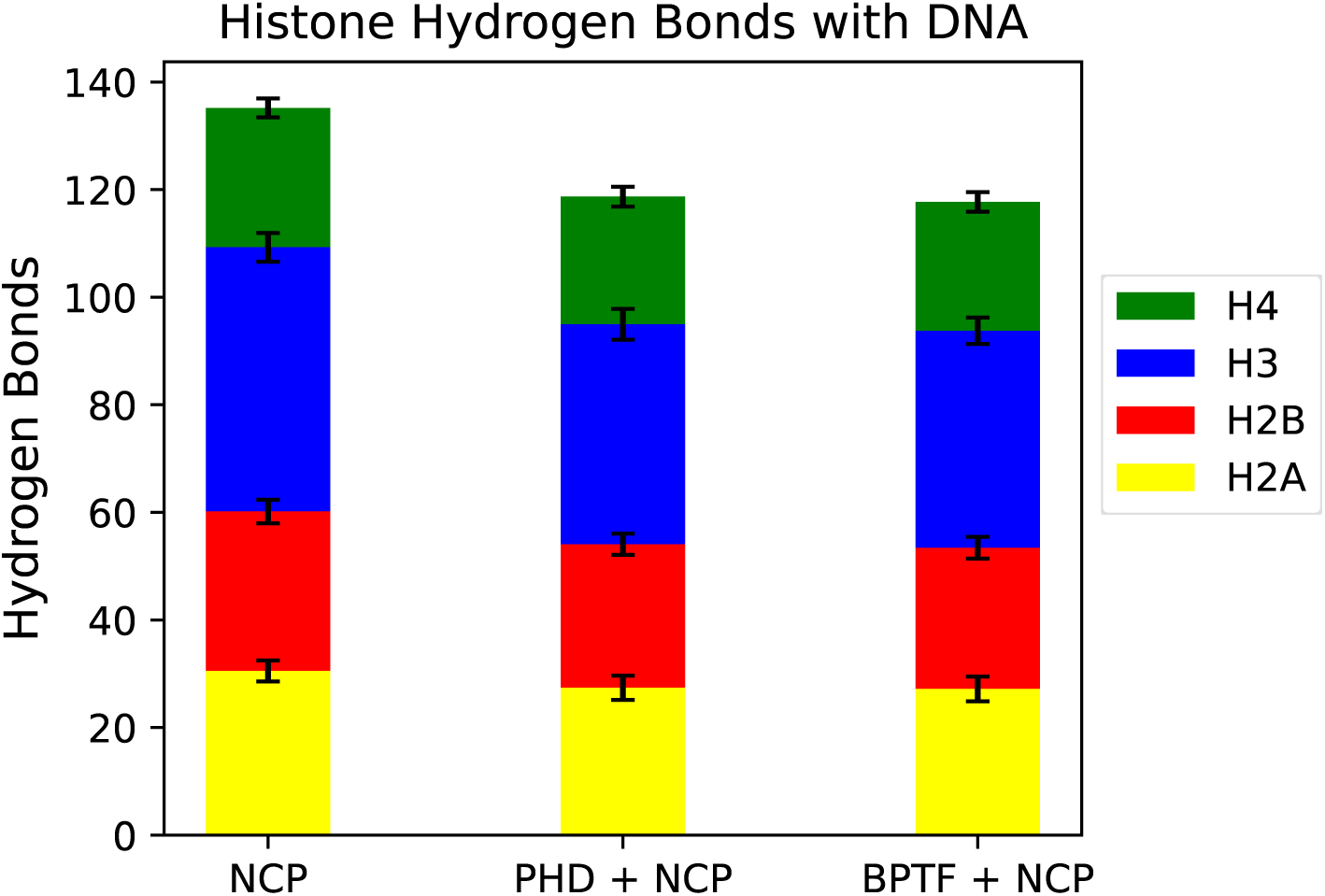
Comparison of the total number of hydrogen bonds between the histones and DNA. This difference is spread across the cores and tails of all of the histones, with both H3 histones constituting a third of this difference despite the PHD finger only being bound to one of them.

**Figure 7:**
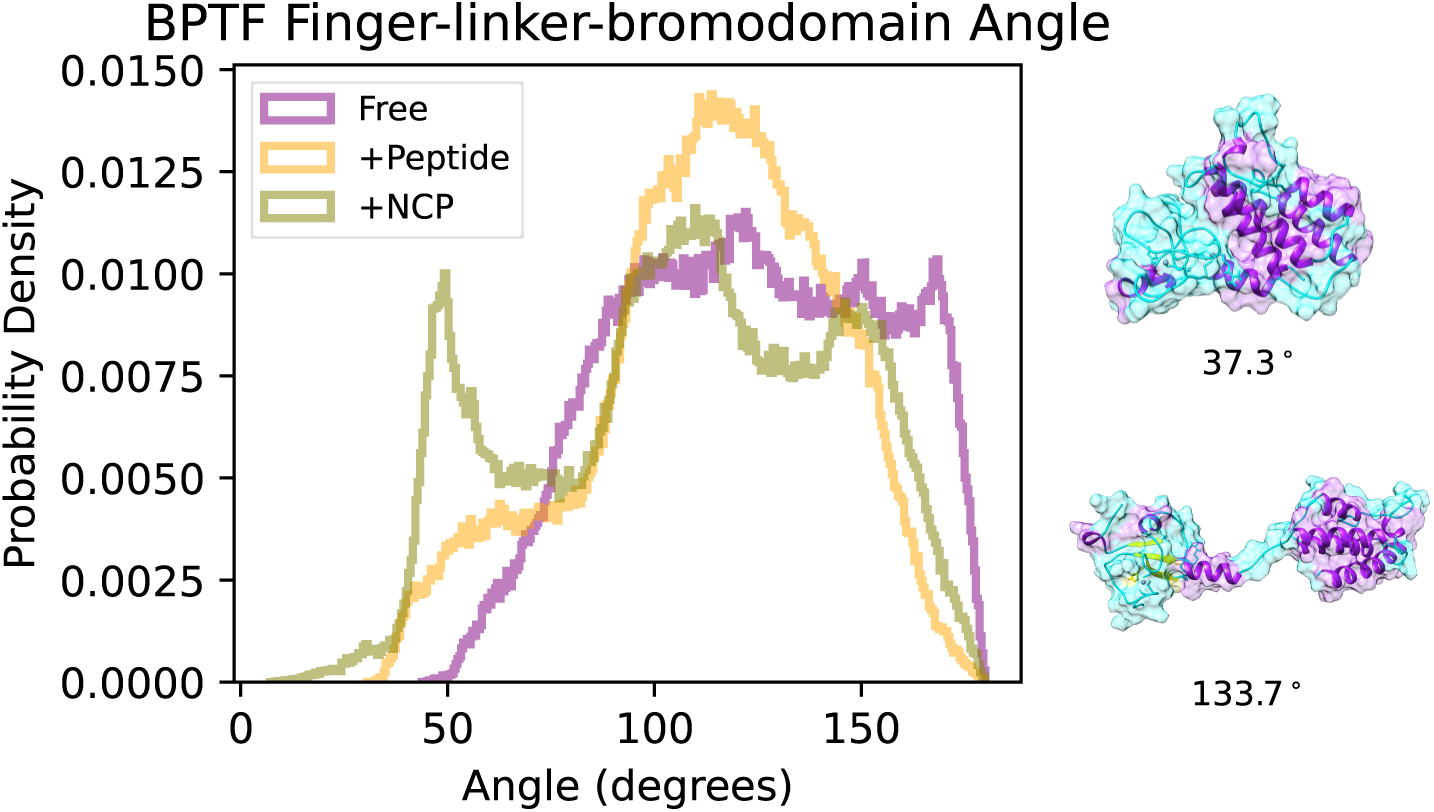
The angle between the bromodomain and the PHD finger-linker axis shows flexibility in the BPTF fragment as the linker between the PHD finger and bromodomain acts as a hinge. H3 binding increases sampling of more compact conformations, with even larger sampling of compact conformations in NCP bound systems. Two sample conformations show the large scale motions of the bromodomain relative to the PHD finger.

Reductions in hydrogen bonds were also observed in other histones. H4–DNA hydrogen bonds dropped from 26±2 in the unbound nucleosome by 2 with both readers. H2A dropped from 31±2 by 3 with both readers, and H2B dropped from 30±2 by 3. Figure 6 summarizes these changes. Overall, reader binding reduced the total number of histone–DNA hydrogen bonds by 16 for the PHD finger and by 17 for BPTF. This represents a significant weakening of core–DNA interactions and supports a model where reader binding loosens the nucleosome structure by disrupting interactions across the core.

### 3.4 Electrostatic Repulsion and Van der Waals Attraction Govern BPTF-Nucleosome Binding

Although the H3 tails collapsed onto the nucleosome core in all simulations, the bound reader domains made little to no direct DNA contact. In reader-bound systems, the PHD domain consistently maintained contact with the H3 tail, which acted as a physical buffer separating the transcription factor from DNA. The larger BPTF fragment was even farther from DNA than the PHD finger.

Despite this limited contact, both readers carry a net negative charge and frequently approached within a Debye length of the negatively charged DNA over the course of the simulations (Figure S16). This consistent proximity indicates that the reader and DNA remained within range of one another for electrostatic interactions throughout most of the post-equilibration period. Further, readers contributed more contacts to H3 while the total number of histone-DNA contacts remained approximately the same.

We performed MM/GBSA analysis to better understand the driving forces behind reader binding. As reader size increased, a smaller fraction of the molecule remained within electrostatic range of the DNA. Consistent with this, PHD-bound systems experienced a stronger electrostatic penalty, with an energy of 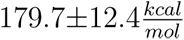, compared to 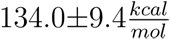 for BPTF-bound systems.

Despite these unfavorable terms, the total binding energy was dominated by highly favorable Van der Waals interactions, which contributed 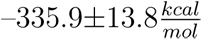 for PHD and 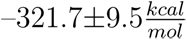 for BPTF (Table 2). This strong short-range attraction matches the increased number of contacts observed between the reader and the H3 tail and offsets the electrostatic cost of approaching the nucleosome. The binding energy terms from the three-trajectory MM/GBSA calculations were larger in magnitude than those from single-trajectory MM/GBSA for the same interactions. Nonetheless, both methods revealed similar trends, with the same interplay between electrostatics and short-range attraction dominating reader binding (Table S5). The three-trajectory approach additionally captured binding-induced conformational changes, including compensatory tail–DNA interactions from other histones and reduced H3–DNA contacts, which are absent from single-trajectory estimates.

**Table 2:** Three trajectory GBSA results for the NCP + Reader systems in kcal/mol. Van der Waals interactions drive the stability of the association between the nucleosome core and reader domains.

| System | $\Delta E_{elec}$ | $\Delta E_{internal}$ | $\Delta E_{VdW}$ | $\Delta E_{total}$ |
| --- | --- | --- | --- | --- |
| PHD + NCP | $179.7 \pm 12.4$ | $23.4 \pm 18.3$ | $-335.9 \pm 13.8$ | $-132.8 \pm 26.1$ |
| BPTF + NCP | $134.0 \pm 9.4$ | $31.8 \pm 14.1$ | $-321.7 \pm 9.5$ | $-155.9 \pm 19.4$ |

These results suggest that electrostatic repulsion between the negatively charged PHD finger and DNA limits direct contact, while strong Van der Waals interactions stabilize the reader’s association with the nucleosome. While MM/GBSA estimates should be interpreted with caution, the relative magnitudes still indicate that short-range dispersive forces are the primary drivers of reader binding. However, the reader appeared to undergo a structural adaptation depending on what it binds. Given that the larger BPTF fragment showed distinct effects on nucleosome dynamics compared to the PHD finger alone, we next examined whether this difference extended to the structural properties of the reader itself.

### 3.5 BPTF Conformations are Altered by Nucleosome Engagement

We observed several differences in nucleosome behavior that depended on reader size, suggesting that the structural and dynamic properties of the readers themselves may play a role. To investigate this, we analyzed how having both the PHD finger and bromodomain influences BPTF’s structure and flexibility, and how these properties change upon nucleosome binding. Regardless of whether BPTF was free, peptide-bound, or nucleosome-bound, our simulations revealed substantial conformational variability between the PHD finger and bromodomain. In all systems, both domains remained relatively stable, with RMSD values relative to the starting conformation for the PHD finger between 1.4 and 7.2 Å and for the bromodomain between 1.3 and 4.9 Å (Figure S17). However, the overall RMSD for the entire BPTF fragment was much higher—up to 15.0 Å—which points to significant motion between domains rather than within them.

To characterize this interdomain flexibility more directly, we measured the angle defined by the PHD finger, linker, and bromodomain. An angle of 0° corresponds to a fully closed conformation, while 180° marks a fully open state. Unbound BPTF sampled angles ranging from 43.8° to 180.0°. Binding lowered this range by 12.9° in the peptide-bound state and by 36.5° in the NCP-bound state, indicating a shift toward more compact conformations as BPTF engaged its binding partners. Time series analysis showed swings of over 100° in all systems, consistent with broad conformational sampling. However, bound BPTF spent more time in compact, closed states, especially when bound to the nucleosome. This increased compaction in the NCP-bound system may underlie the more pronounced effects of BPTF on nucleosome core dynamics compared to the PHD finger alone.

Analysis of the radius of gyration (RG) reinforced these findings. Across all systems, RG ranged from 17.5 to 29.1 Å, and the RG mode shifted depending on binding state. The unbound BPTF was more extended, with an RG mode of 25.0 Å. Peptide binding decreased the RG mode by 1.2 Å, and NCP binding decreased it by 3.1 Å (Figure S18). Although BPTF remained highly flexible in all states, the nucleosome-bound form was more frequently compact, in contrast to the more open conformations sampled by unbound and peptide-bound BPTF.

Binding state also altered the volume and stability of the H3-binding pocket in the PHD finger. In solution, the volume was 1132±49.6 Å^3^ for the PHD finger and 1105±46.7 Å^3^ for BPTF. Binding to the H3 peptide increased the average volume by 90.8 and 114.1 Å^3^ in order of increasing reader size; volumes increased further from unbound to NCP-bound by 115.8 and 145.7 Å^3^ (Figure 8).

**Figure 8:**
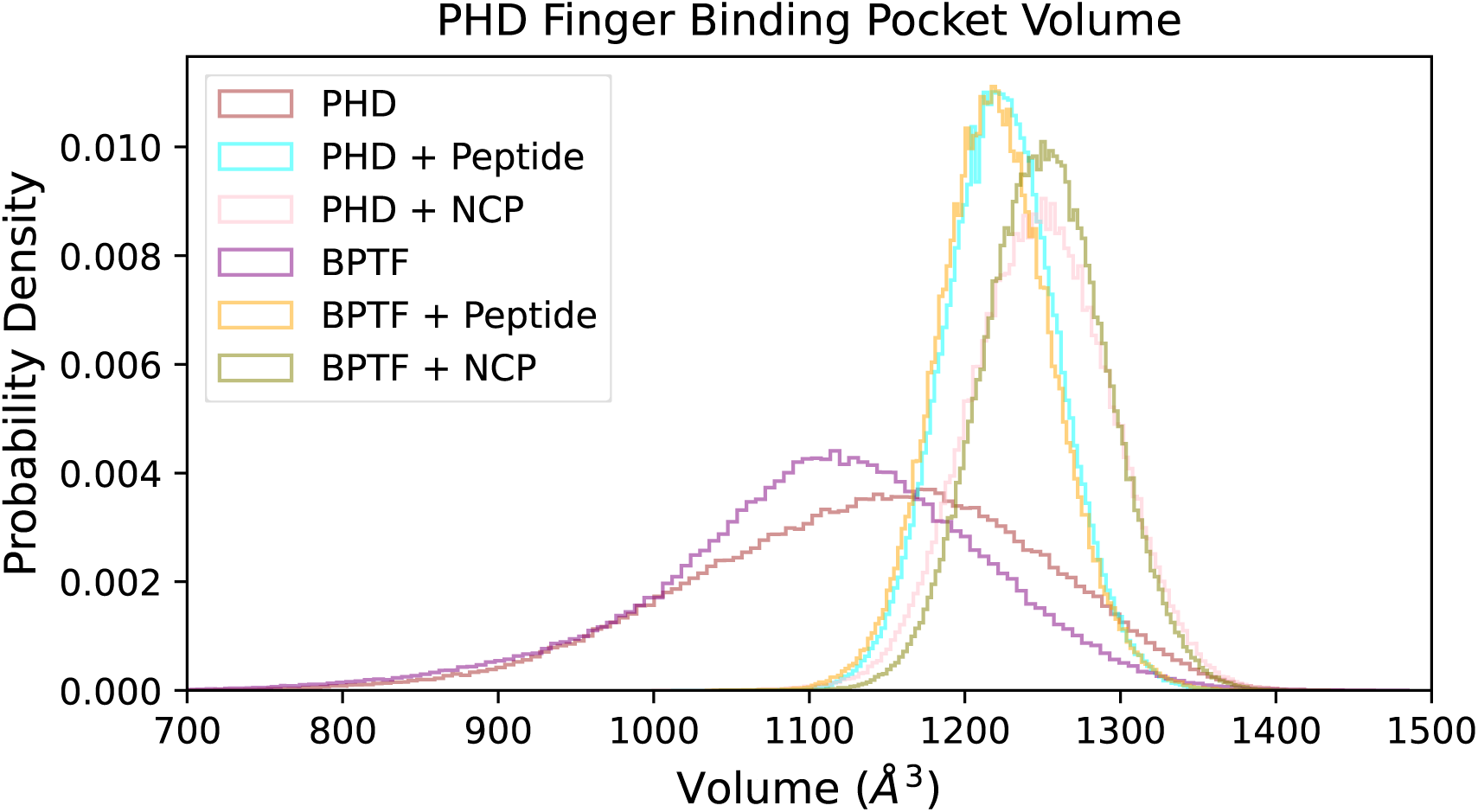
Binding pocket volume of the PHD finger in all systems. H3 binding keeps the pocket open, and nucleosome proximity appears to augment this effect.

This expansion was accompanied by greater stability in the binding pocket region, as reflected in lower RMSF values for heavy atoms in PHD finger residues 17–48 (Figure 9). Peptide binding reduced these RMSFs by 0.4 to 0.5 Å, while NCP binding reduced them by 0.2 to 0.3 Å. Over time, unbound systems exhibited larger pocket volume fluctuations than either bound system (Figure S19). These results indicate that H3 binding both stabilizes the PHD finger and holds the binding pocket open in peptide- and NCP-bound systems, with a slightly greater opening observed in the NCP-bound state.

**Figure 9:**
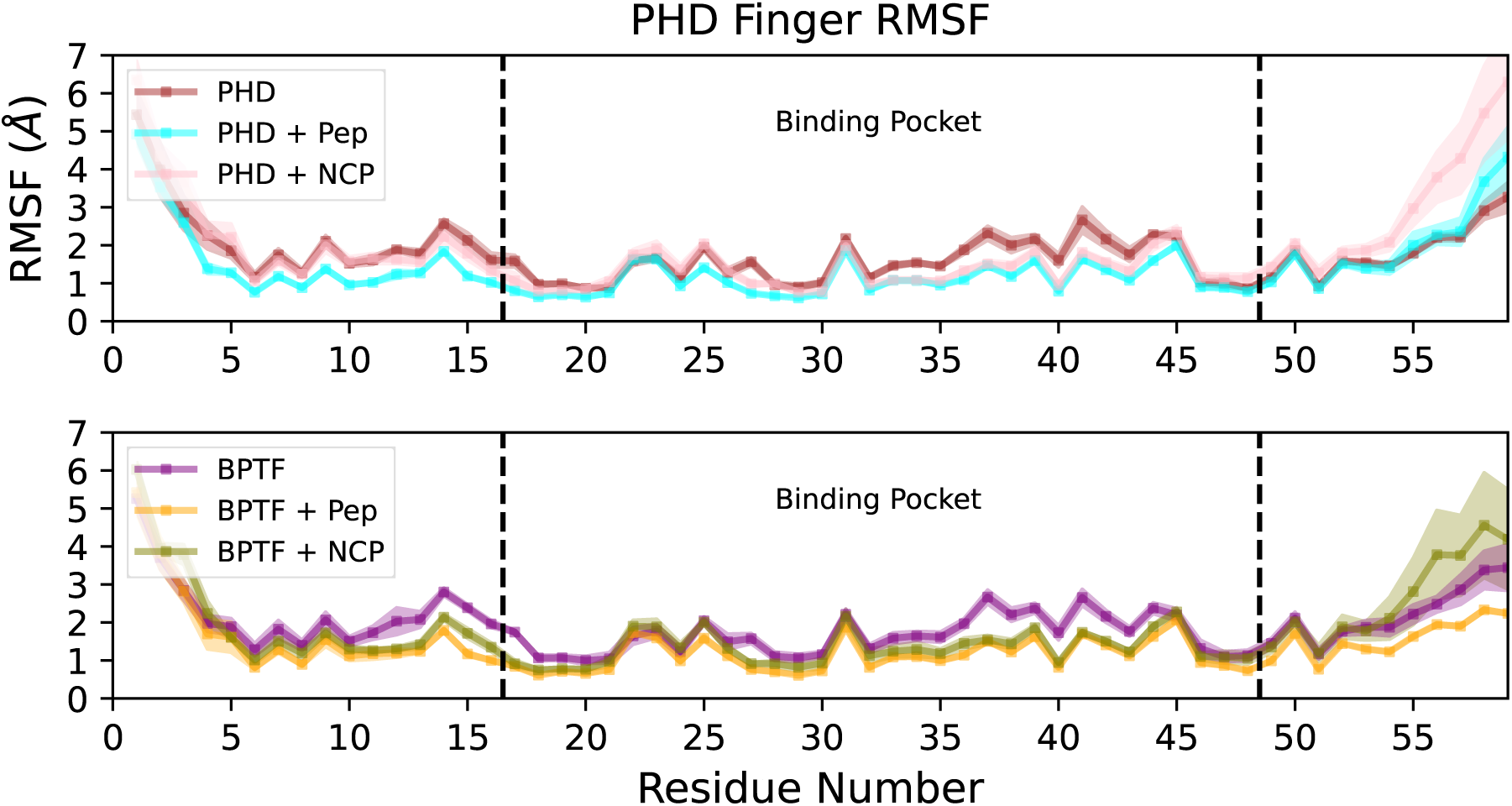
Top: Root mean square fluctuations of the first 59 PHD finger residues from all systems. H3 binding stabilizes the binding pocket region, with a smaller effect on the full-length nucleosome.

Taken together, these findings show that BPTF remains flexible but does not adopt a single static conformation. Instead, its multidomain structure adapts to the binding partner, supporting stable engagement with the nucleosome. This structural adaptation fits with the favorable Van der Waals interactions observed in our MM/GBSA analysis and may help BPTF overcome electrostatic penalties to exert a broader impact on nucleosome dynamics than the PHD finger alone.

## 4 Discussion

Our study used all-atom MD simulations to explore how the binding of PTM reader domains, including multidomain constructs like BPTF, influences nucleosome dynamics. PTMs such as H3K4me3 reduce histone tail association with DNA,^30^ exposing sites for reader engagement, including the PHD finger binding pocket examined in this study. In our simulations, the PHD finger began bound to H3K4me3 in an extended tail conformation, allowing the tail to collapse toward the core during equilibration. Systems containing full-length H3 or the H3 1–15 peptide showed greater reader stability than peptide-free systems. Analyses of PHD binding pocket volume, RMSF, RMSD, and BPTF radius of gyration (Figures 8, 9, S17 & S18) revealed distinct conformational dynamics for BPTF when bound to the full nucleosome compared to a short peptide. Competition between the PHD finger and DNA for access to the H3 tail (Figure 2) and MM/GBSA results (Table 2) suggest that DNA remains a relevant binding competitor, even after PHD engagement. The persistence of this competition, despite trimethylation not altering lysine charge, highlights the importance of non-electrostatic contributions to H3 tail positioning.

Despite competition for the H3 tail, the PHD finger made minimal direct contact with DNA. In simulations that included the bromodomain, this domain did not engage the nucleosome core or any histone tails. Across all runs, it consistently adopted an orientation facing away from the NCP toward the solvent. This solvent-facing orientation suggests an entropic penalty for associating with H4 within the same nucleosome as the PHD-bound H3. It may also indicate that bromodomain association following PHD engagement requires additional nucleosomal PTMs or cofactor proteins that were absent from our model systems.

PHD fingers frequently interact with other proteins while simultaneously binding to H3.^86,87^ In our simulations, electrostatic interactions with DNA, increased H3 tail fluctuations, and variability in tail-reader positioning on the nucleosome suggest that PHD–H3 interactions are not strictly constrained by DNA-binding specificity. This observation is consistent with the diversity of chromatin functions associated with PHD fingers. PHD fingers that recognize H3K4me3 occur in a wide variety of proteins involved in transcriptional activation, repression, and remodeling,^45,88–91^ implying that the larger complex determines the functional outcome of H3 engagement. Based on this, the differences in histone and DNA hydrogen bonding observed between reader bound and unbound NCP systems may reflect an early structural change that facilitates recruitment or activity of chromatin remodelers such as hNURF. If these changes are a general feature of PHD finger binding, additional proteins in complex could reinforce or counteract the resulting destabilization to tune DNA accessibility.

We made no other changes to the amino acid or nucleic acid sequences beyond remodeling the H3 tails, which likely preserved the superhelical positions of other histone tails from the crystal structure. The increased H4–DNA binding we observed may reflect these starting conformations combined with reader-induced DNA exposure near the bound H3, rather than a direct effect on H4 itself. This observation aligns with the behavior of the BPTF fragment, where flexibility between the PHD finger and bromodomain may influence how it engages chromatin. Cooperative effects between these domains have been reported in other finger–linker–bromodomain systems, independent of linker length.^89^ In our simulations, BPTF consistently adopted more compact conformations when bound to the full nucleosome compared to the H3 fragment, indicating a preference for intramolecular interactions in a full chromatin context. Although we did not observe simultaneous engagement of both the PHD finger and bromodomain, the loop connecting them remained flexible, and additional components of the full BPTF complex may restrict this motion. The bromodomain consistently oriented away from the nucleosome, reducing the likelihood of direct interaction. Our nucleosomes lacked H4 acetylation marks, which are important for bromodomain binding,^39^ particularly when combined with H3K4me3 recognition by the PHD finger. Increased H4–DNA association following PHD binding likely further limits bromodomain access in mononucleosome systems.

At the sampled time scales, we observed increased H3 tail fluctuations resulting from both trimethylation and reader binding. BPTF-containing complexes induce nucleosome sliding, a remodeling feature that occurs on timescales not accessible to conventional all-atom simulations,^92^ although it has been captured in coarse-grained models.^93^ This study used a mononucleosome, yet the initial association of the reader with the histone tail likely depends on intermolecular forces present *in vivo* but absent in these simplified systems.^30,31^ Multiple coexisting PTMs are expected within a given histone octamer,^94^ and different combinations could drive diverse cross-talk effects. These may include additional H3 modifications that reduce PHD finger binding affinity and H4K16ac, which enables BPTF’s canonical binding mode.^39^ In our simulations, we observed a reduction of 16–17 hydrogen bonds between histones and DNA upon PHD finger or BPTF binding. This reduction is comparable to the total number of H3 tail–DNA hydrogen bonds, suggesting that even limited reader engagement can significantly weaken core histone–DNA interactions (Table S1). While we observed localized shifts in histone–DNA contacts, large-scale unwrapping or nucleosome destabilization did not occur on this timescale. Altogether, our results show how BPTF reader engagement modulates nucleosome structure through both direct H3 tail interactions and propagated effects on histone dynamics, providing insight into the allosteric influence of reader domains within chromatin.

## 5 Data availability

All simulation inputs, scripts used for analysis, and processed data files are available at https://github.com/WereszczynskiGroup/supplemental-data-hebert-bptf-2025. Raw trajectory data generated in this study are available on Zenodo at https://doi.org/10.5281/zenodo.14926478. These trajectories are water-stripped and temporally strided to reduce file size but cover the full duration of each simulation. Together, these resources are sufficient to reproduce all analyses and key results reported in the manuscript.

## 6 Declaration of generative AI and AI-assisted technologies in the writing process

During the preparation of this work the authors used ChatGPT in order to refine the manuscript language. After using this tool/service, the authors reviewed and edited the content as needed and take full responsibility for the content of the publication.

## 7 Author Contributions

R.H. prepared and executed the simulations, performed the analyses, and generated the figures presented in this paper. R.H. and J.W. designed the study, interpreted the results, wrote the manuscript, and edited and approved the final version.

## 8 Acknowledgments

This project was supported by the National Institutes of Health grant R35GM119647.

