## Supporting Information for "Deciphering the Molecular Mechanisms of BPTF Interactions with Nucleosomes via Molecular Simulations"

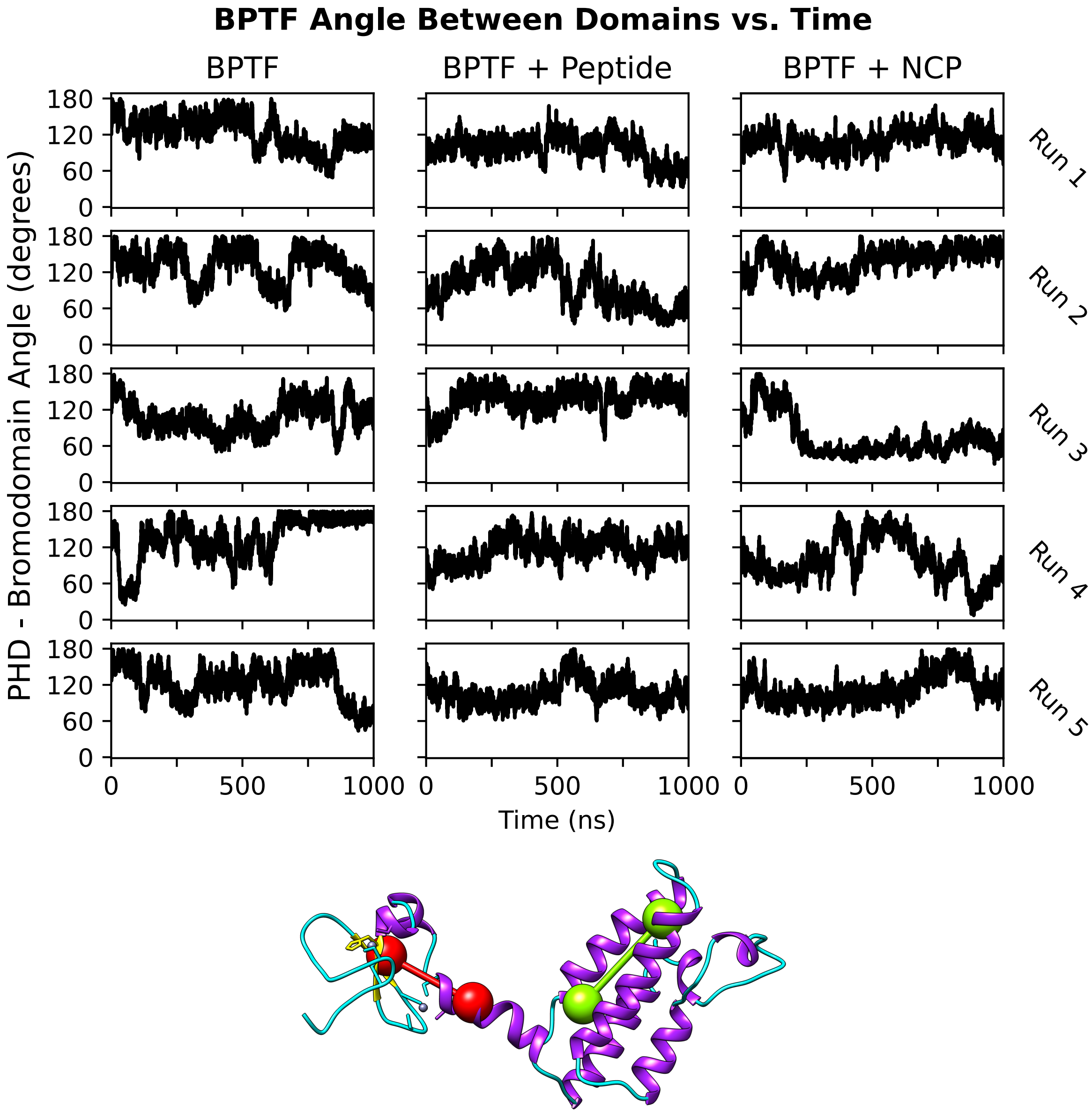


Figure S1: Angle vs. time between the two stable domains of BPTF by system and simulation. The angle is formed by axes along the PHD finger and helical linker centroids (red) and between two helix centroids in the bromodomain (green). Most systems sample an opening and closing motion along a large range of angles, with all systems experiencing fully open conformations in at least one simulation and large variation across simulations. The largest and quickest variation in individual simulations generally occurs in the absence of any other bound protein. NCP and peptide binding appear to generally slow this variation but sampling indicates large stochasticity in this inter-domain angle.

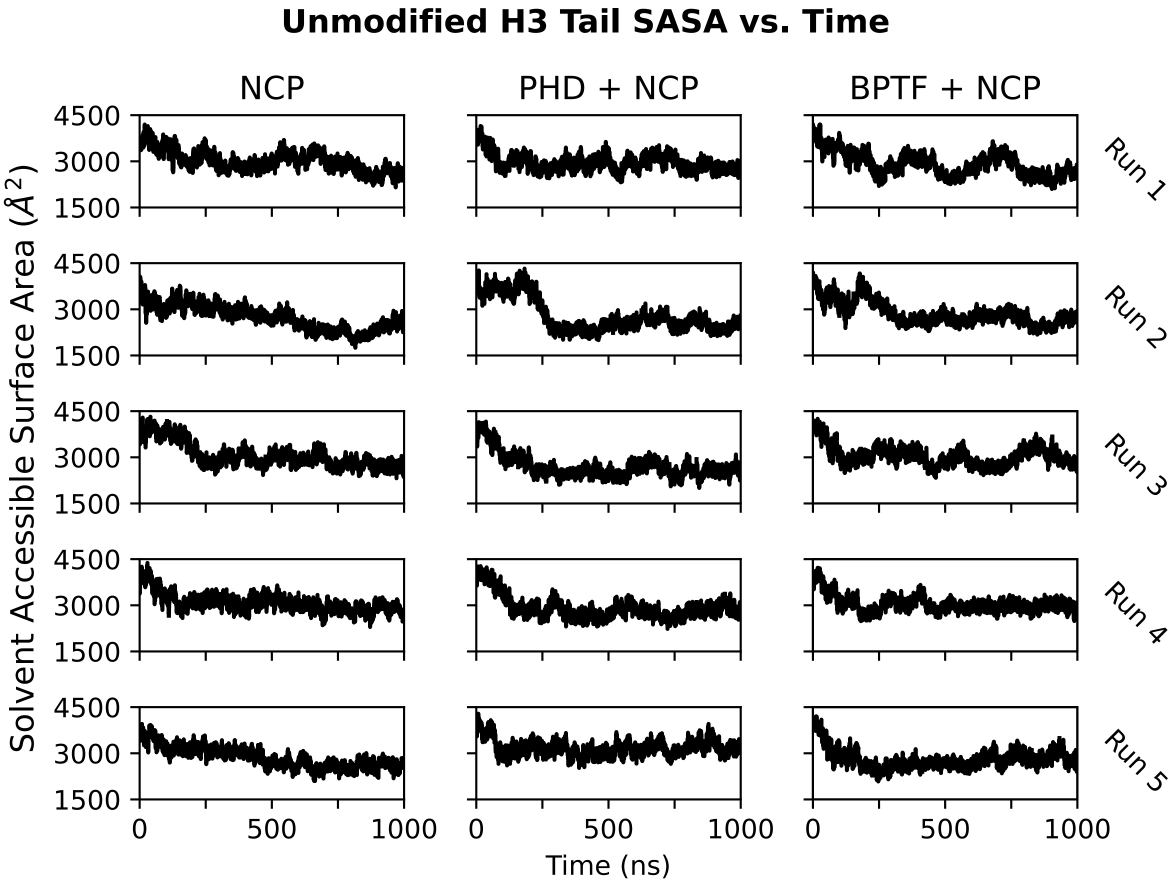


Figure S2: Solvent accessible surface area (SASA) of the unmodified H3 tail over time. H3 tails started extended into solution and were allowed to collapse onto the core. Associating with DNA results in a reduction in SASA.

Table S1: Single trajectory MM/GBSA results for the modified H3 tail - DNA interactions in kcal/mol. Reader binding led to a reduction in Van der Waals interactions betrween H3 and DNA, reducing the overall binding affinity.

| System | $\Delta E_{elec}$ | $\Delta E_{VdW}$ | $\Delta E_{total}$ |
|---|---|---|---|
| NCP | $3.2 \pm 4.4$ | $-67.8 \pm 3.0$ | $-64.6 \pm 5.3$ |
| PHD + NCP | $6.6 \pm 10.5$ | $-52.0 \pm 6.9$ | $-45.4 \pm 12.6$ |
| BPTF + NCP | $8.2 \pm 5.7$ | $-48.8 \pm 3.7$ | $-40.6 \pm 6.8$ |

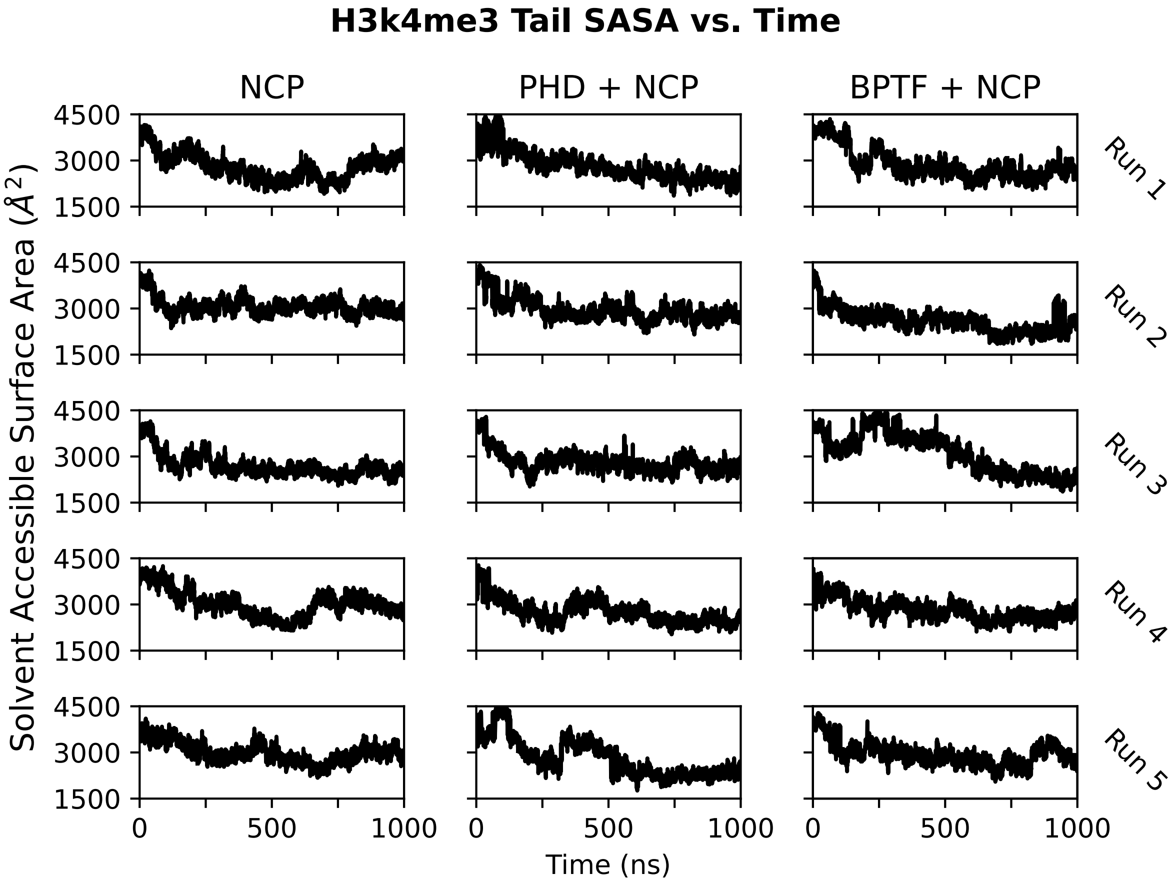


Figure S3: SASA of the modified H3 tail over time. These tails also start simulations extended and collapse onto the core. Associating with DNA and the reader reduces SASA.

Table S2: Single trajectory MM/GBSA results for the unmodified, unbound H3 tail - DNA interactions in kcal/mol.

| System | $\Delta E_{elec}$ | $\Delta E_{VdW}$ | $\Delta E_{total}$ |
|---|---|---|---|
| NCP | $12.2 \pm 5.3$ | $-88.4 \pm 3.2$ | $-76.2 \pm 6.2$ |
| PHD + NCP | $13.7 \pm 7.2$ | $-82.1 \pm 4.7$ | $-68.5 \pm 8.6$ |
| BPTF + NCP | $11.3 \pm 3.5$ | $-88.9 \pm 2.2$ | $-77.6 \pm 4.1$ |

Table S3: Single trajectory MM/GBSA results for H4 tail - DNA interactions in kcal/mol near the modified H3. Reader binding of H3 appears to increase Van der Waals interactions of nearby H4 with DNA.

| System | $\Delta E_{elec}$ | $\Delta E_{VdW}$ | $\Delta E_{total}$ |
|---|---|---|---|
| NCP | $-1.3 \pm 10.0$ | $-38.0 \pm 6.4$ | $-39.3 \pm 11.9$ |
| PHD + NCP | $6.8 \pm 7.0$ | $-64.7 \pm 4.9$ | $-57.9 \pm 8.5$ |
| BPTF + NCP | $6.6 \pm 9.9$ | $-58.7 \pm 6.0$ | $-52.1 \pm 11.6$ |

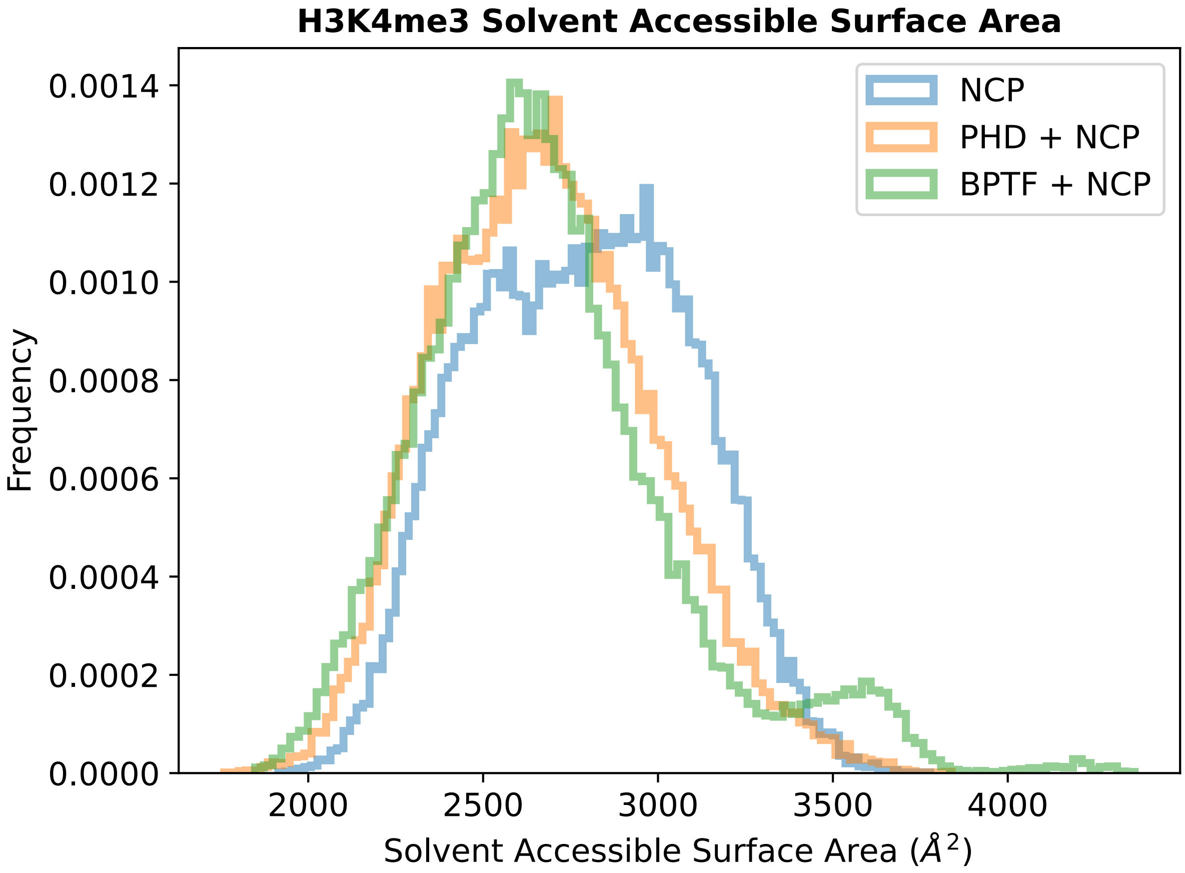


Figure S4: Post-equilibrated calculation of the solvent accessible surface area (SASA) of the modified H3 tail. Reader binding reduces the H3 tail's SASA because the tail tends to settle associating with both DNA and the reader.

Table S4: Single trajectory MM/GBSA results for H4 tail - DNA interactions in kcal/mol near the unmodified H3. There is no apparent effect from reader binding of the modified H3 for this tail, and energy values are similar to those of H4 near bound H3.

| System | $\Delta E_{elec}$ | $\Delta E_{VdW}$ | $\Delta E_{total}$ |
|---|---|---|---|
| NCP | $8.3 \pm 4.1$ | $-61.1 \pm 2.5$ | $-52.8 \pm 4.8$ |
| PHD + NCP | $14.2 \pm 6.4$ | $-71.0 \pm 4.3$ | $-56.8 \pm 7.7$ |
| BPTF + NCP | $10.8 \pm 2.5$ | $-64.4 \pm 1.5$ | $-53.7 \pm 2.9$ |

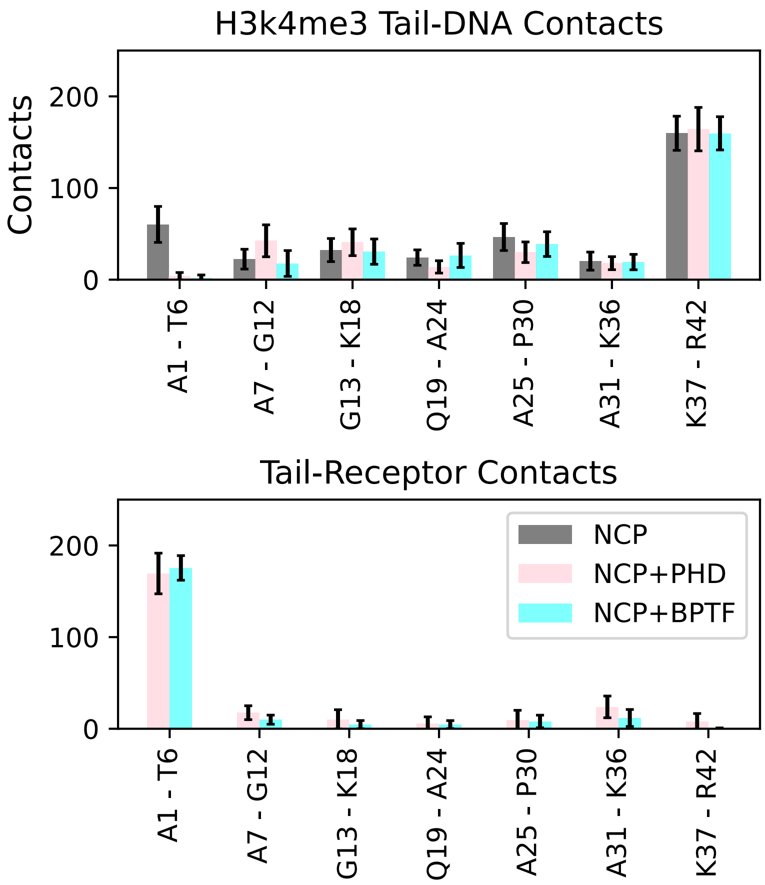


| | H3 1-6 DNA Contacts | H3 1-6 Reader Contacts |
|---|---|---|
| NCP | 60 ± 20 | -- |
| PHD + Peptide | -- | 181 ± 10 |
| NCP + PHD | 4 ± 4 | 169 ± 22 |
| BPTF + Peptide | -- | 182 ± 10 |
| NCP + BPTF | 2 ± 3 | 175 ± 13 |

Figure S5: The distribution of contacts over the first 42 residues of the modified H3 tail, in batches of 6. Top: H3 tail contacts with DNA in all NCP systems. Middle: Tail contacts with the reader protein in NCP systems. Bottom: A table summarizing the split in contacts between the reader and DNA for the first 6 H3 residues.

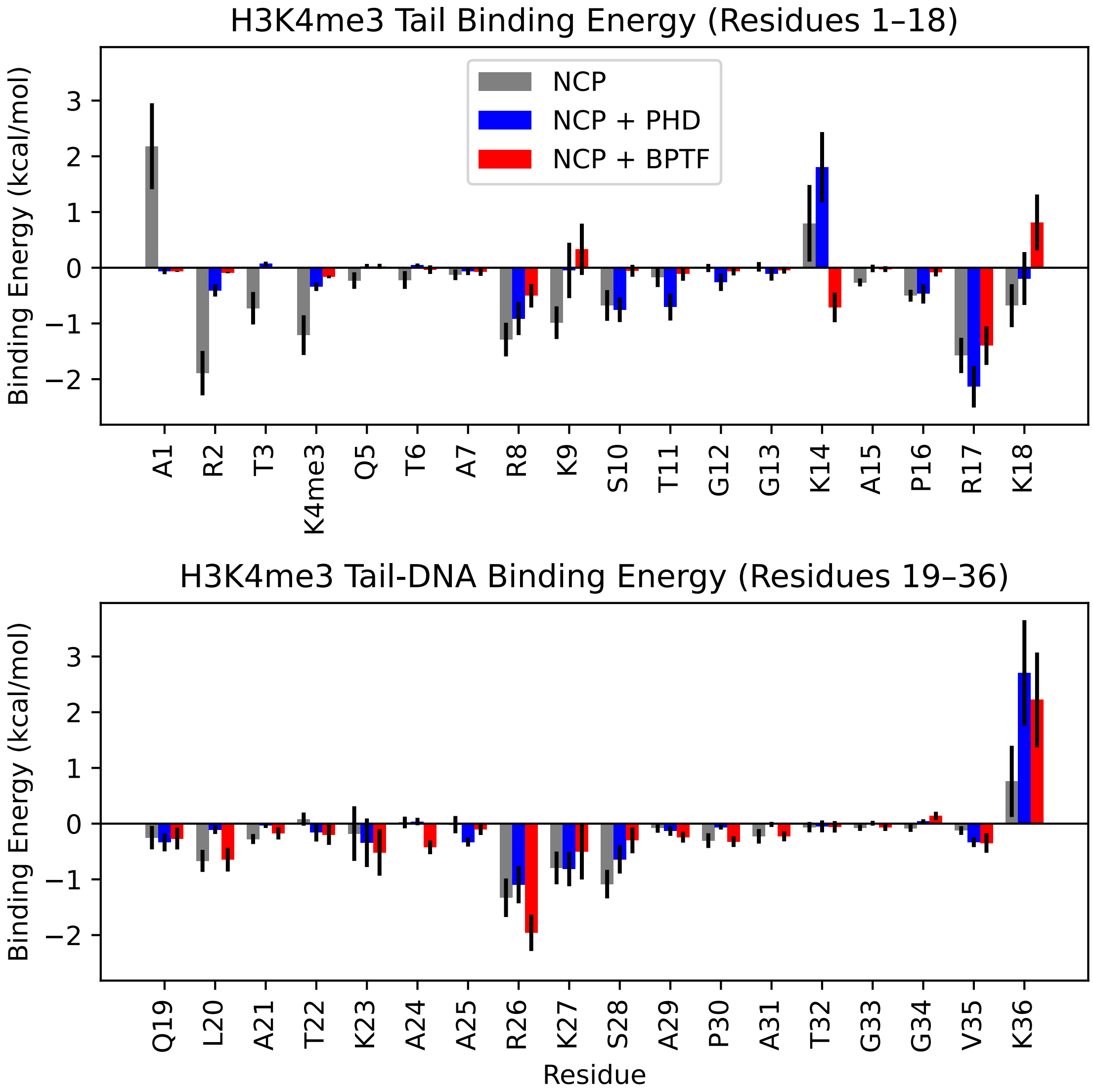


Figure S6: MM/GBSA residue decomposition of the H3K4me3 tail in tail - DNA interactions. Reader binding knocked out most DNA interactions for the first 7 residues.

Table S5: Single trajectory MM/GBSA results for reader - DNA interactions in kcal/mol. There is no apparent effect from reader binding of the modified H3 for this tail, and energy values are similar to those of H4 near bound H3.

| System | $\Delta E_{elec}$ | $\Delta E_{VdW}$ | $\Delta E_{total}$ |
|---|---|---|---|
| PHD + NCP | $42.2 \pm -25.4$ | $-94.9 \pm 20.2$ | $-52.7 \pm 32.5$ |
| BPTF + NCP | $32.0 \pm 25.4$ | $-75.8 \pm 15.4$ | $-43.8 \pm 29.7$ |

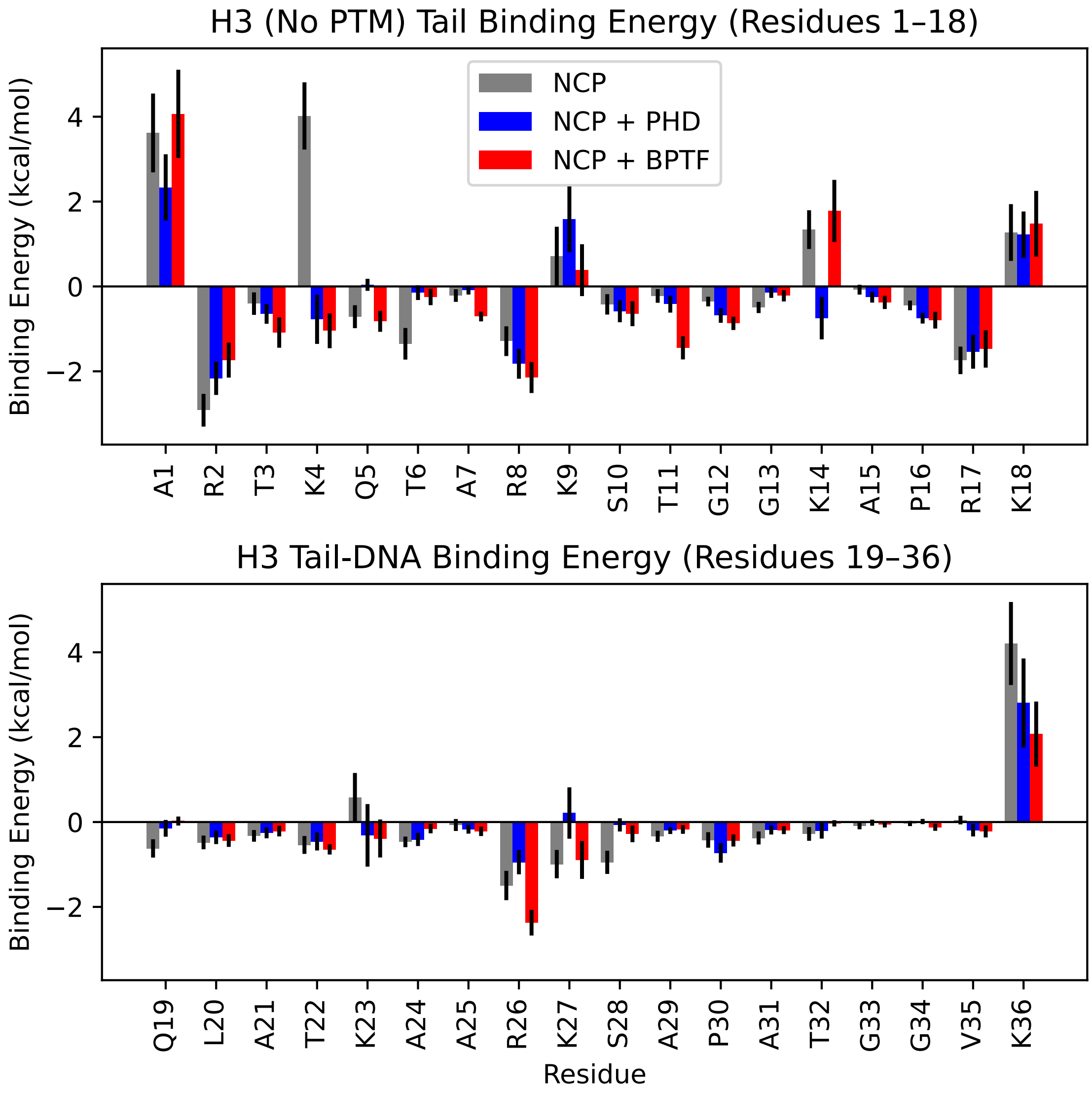


Figure S7: MM/GBSA residue decomposition of the unmodified tail in tail - DNA interactions. The first 7 residues contribute to the interaction, unlike on the bound tail. More of the tail actively contributes to the overall binding energy.

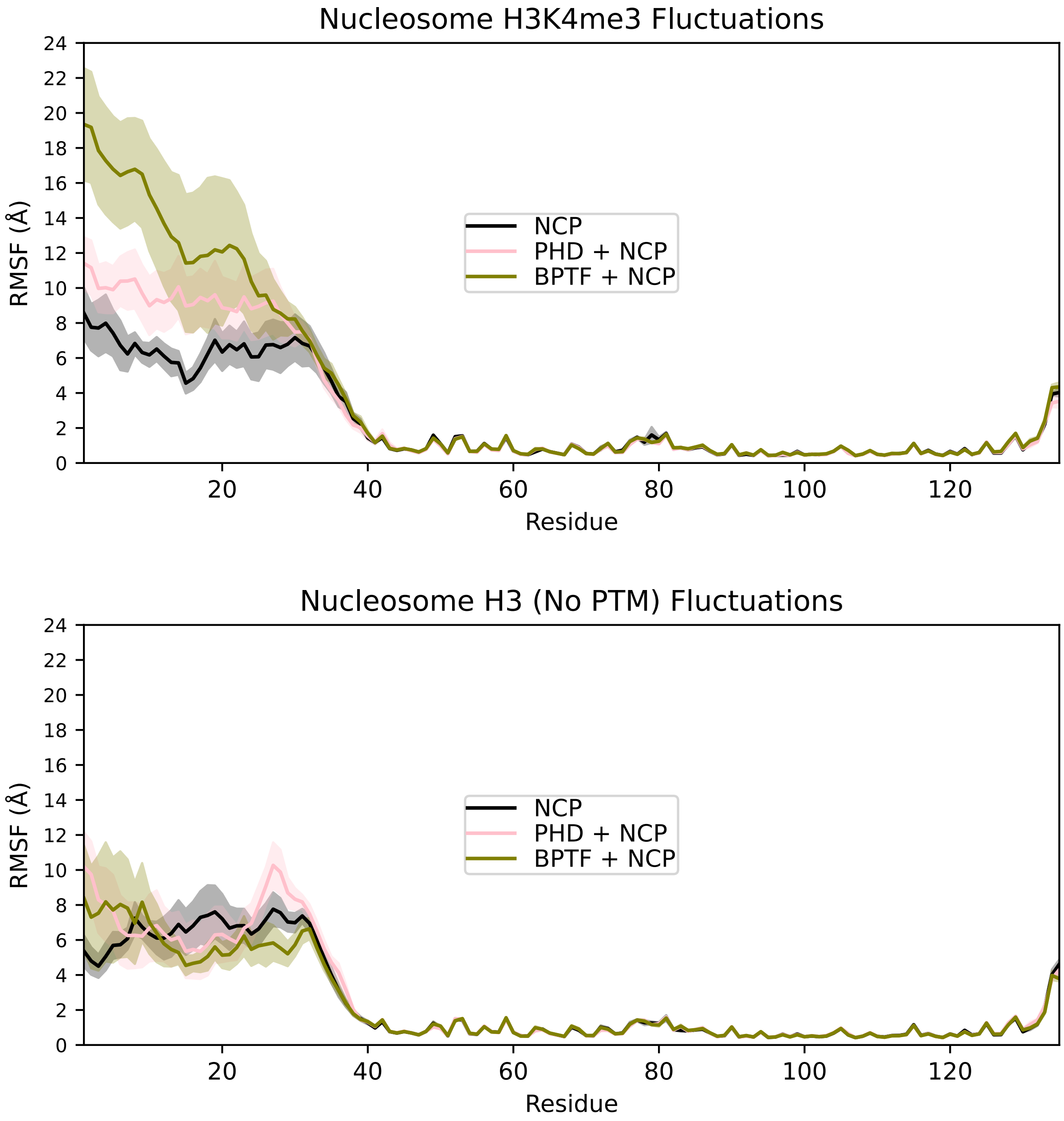


Figure S8: Fluctuations of H3 proteins. Top: Reader binding results in increased fluctuations of the bound H3 tail. A significantly larger increase occurs in BPTF bound systems. Bottom: Adjacent H3.

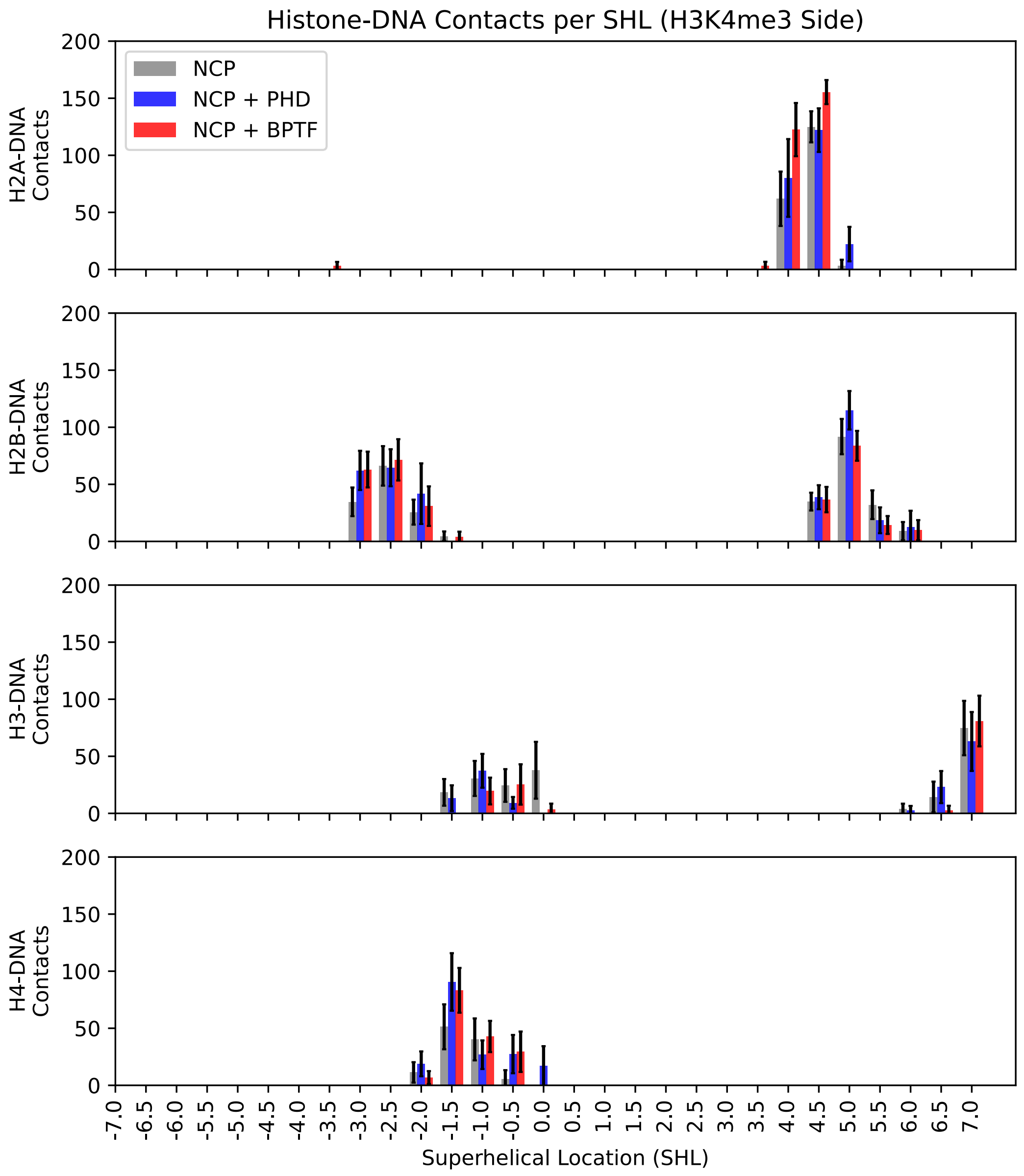


Figure S9: Contacts of histone tails by superhelical location on the same side of the nucleosome dyad as the PTM. SHLs -1.5 to 0.0 have substantial overlap between the two histone tails. SHLs -1.5, -0.5, and 0.0 show correspondence between decreased H3 tail contacts and increased H4 tail contacts from reader binding.

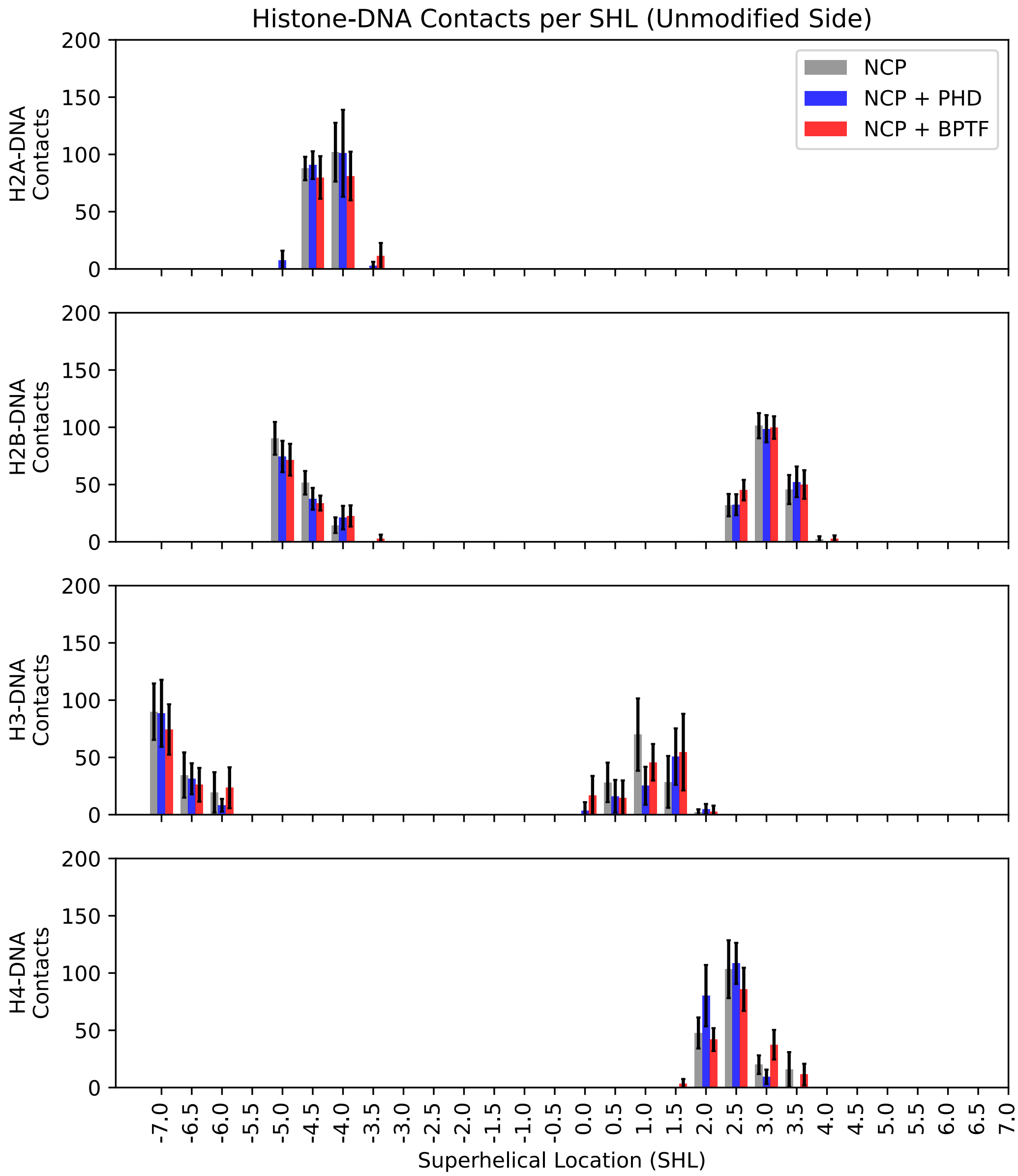


Figure S10: Contacts of histone tails by superhelical location on the opposite side of the nucleosome dyad as the PTM. Overlap in binding location is seen in H2A with H2B tails, as well as H2B with H4 tails.

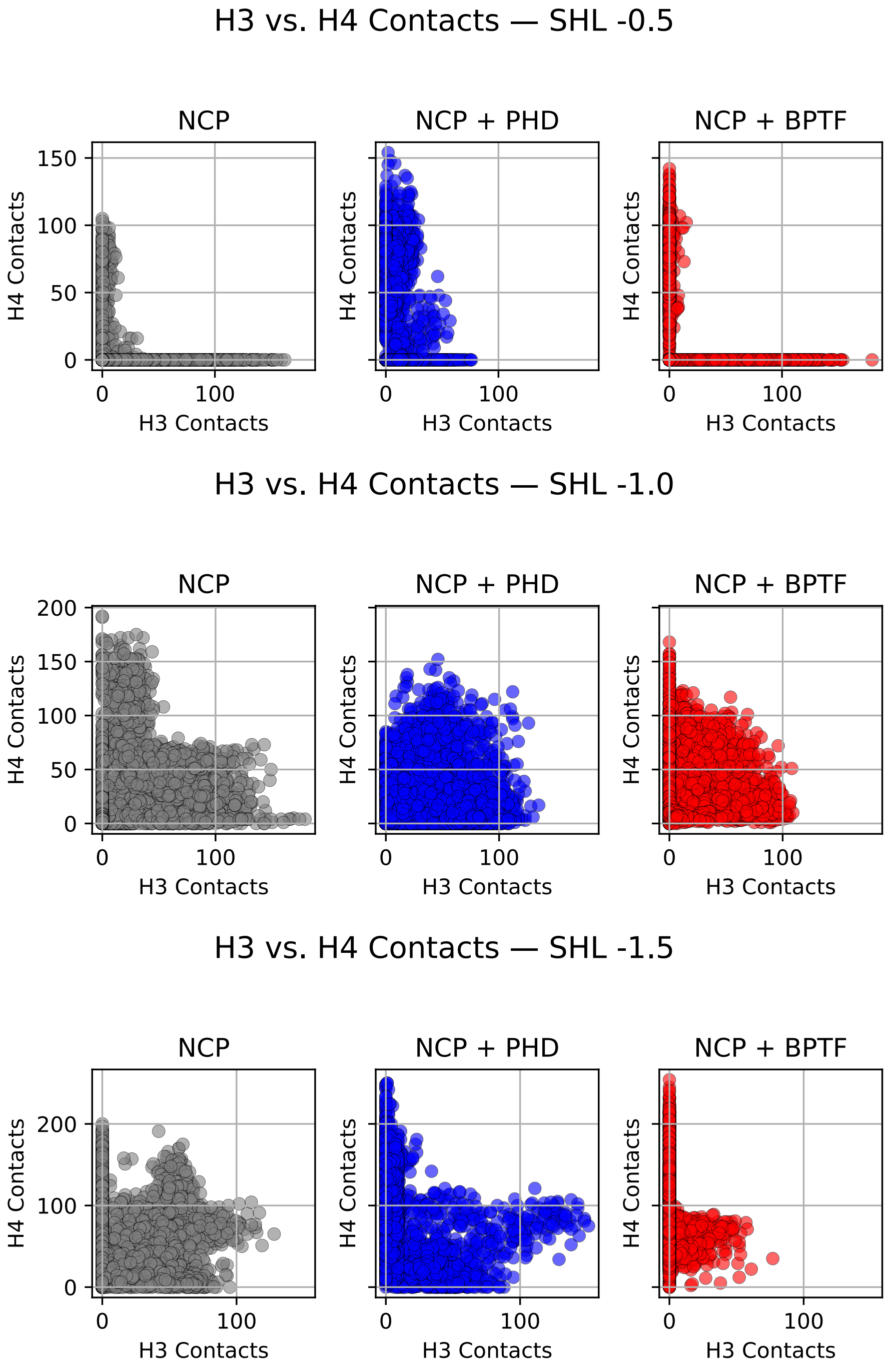


Figure S11: H3 and H4 tails make simultaneous contact with DNA in superhelical locations -0.5 to -1.5, five to fifteen base pairs from the dyad. Simultaneous contacts indicate potential competition for DNA binding.

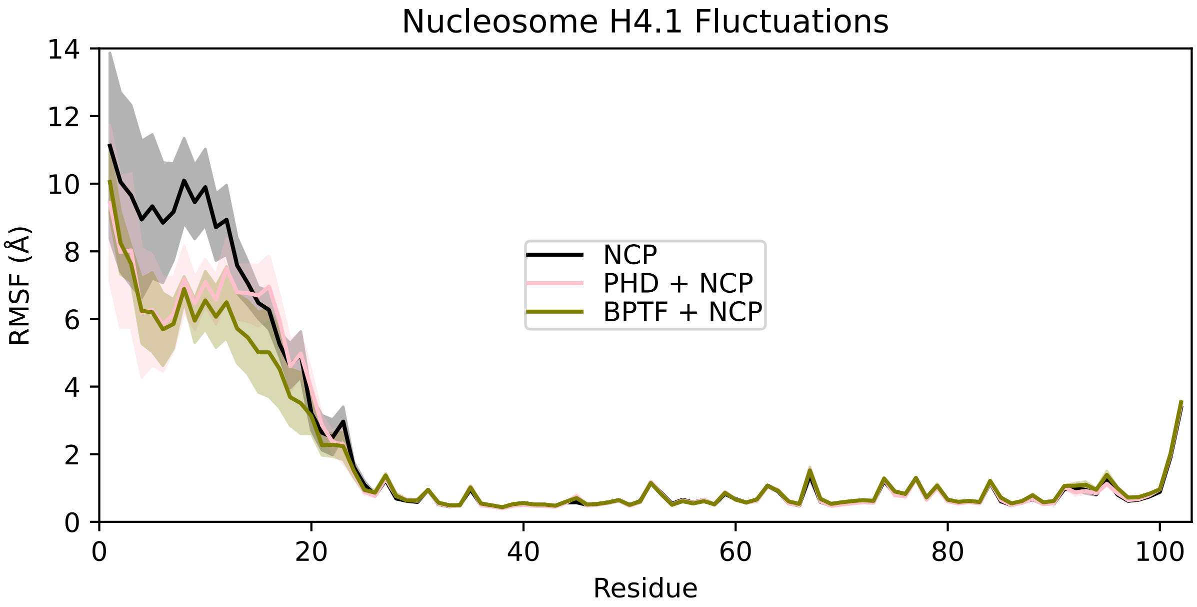


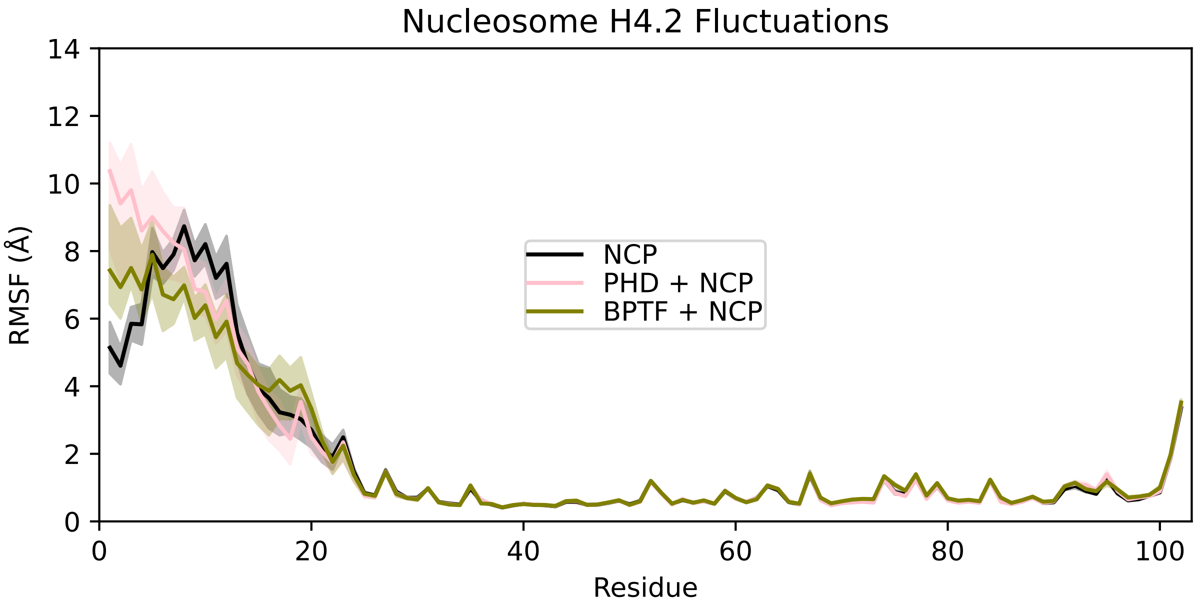


Figure S12: Fluctuations of H4 proteins. Top: H4 on the same side of the dyad axis as the modified H3. Much of the tail sees decreased fluctuations after reader binding. Bottom: H4 on the opposite side of the dyad axis from the modified H3.

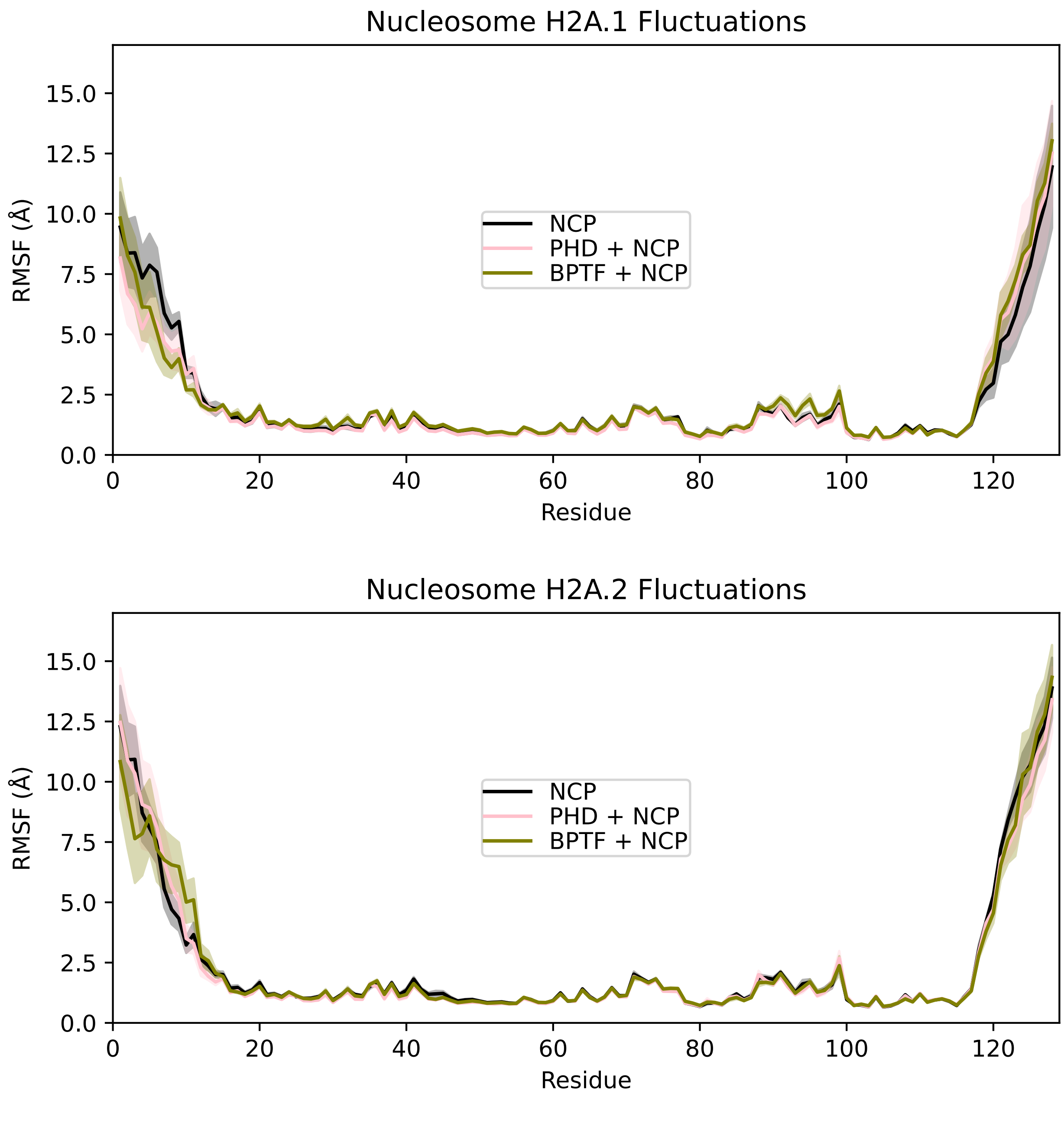


Figure S13: Fluctuations of H2A proteins. Top: H2A on the same side of the dyad axis as the modified H3. Bottom: H2A on the opposite side of the dyad axis from the modified H3.

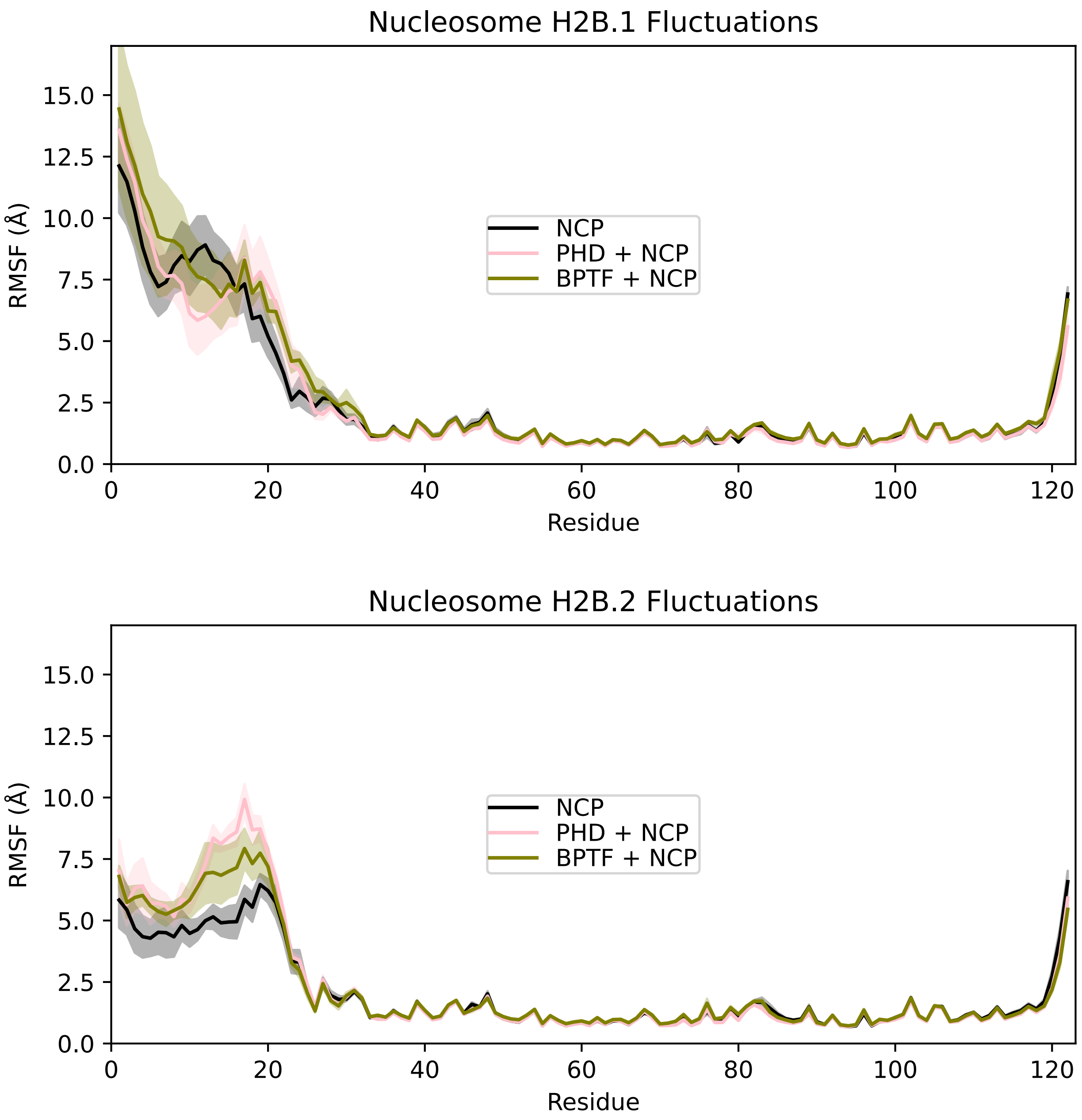


Figure S14: Fluctuations of H2B proteins. Top: H2B on the same side of the dyad axis as the modified H3. Bottom: H2B on the opposite side of the dyad axis from the modified H3.

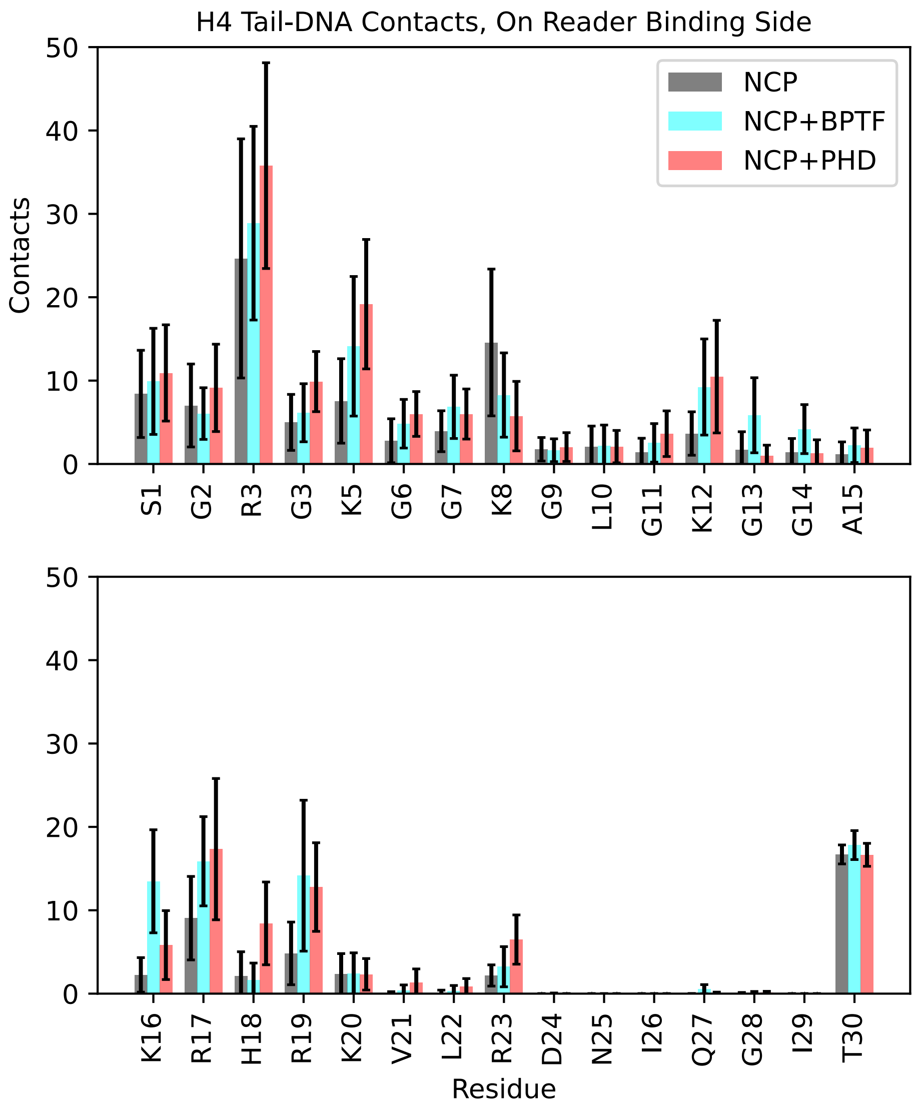


Figure S15: Contacts by residue of histone H4's tail with DNA on the reader-bound side of the NCP. Binding H3 with the reader results in increased H4 contacts with DNA in almost every tail residue, which collectively result in a significant increase in total H4-DNA contacts.

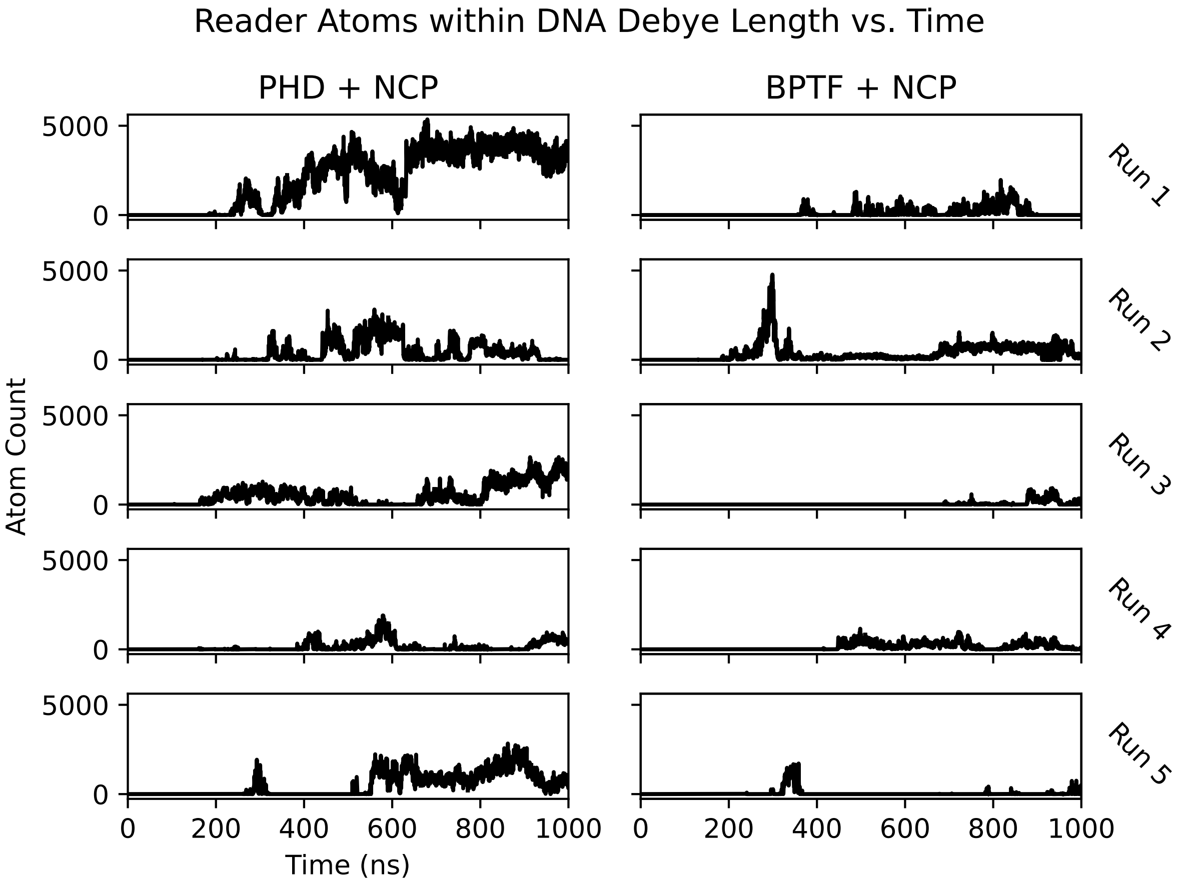


Figure S16: Time series of the atom counts of the PHD finger and BPTF within a Debye length of DNA throughout the simulations. Nonzero values indicate atoms in the protein are subject to the electrostatic field of DNA through solution. Both readers possess electronegative charge, leading to an electrostatic repulsion from DNA within this distance. BPTF spends less time within this distance, while the PHD finger spends more time in this range, leading to a stronger electrostatic repulsion.

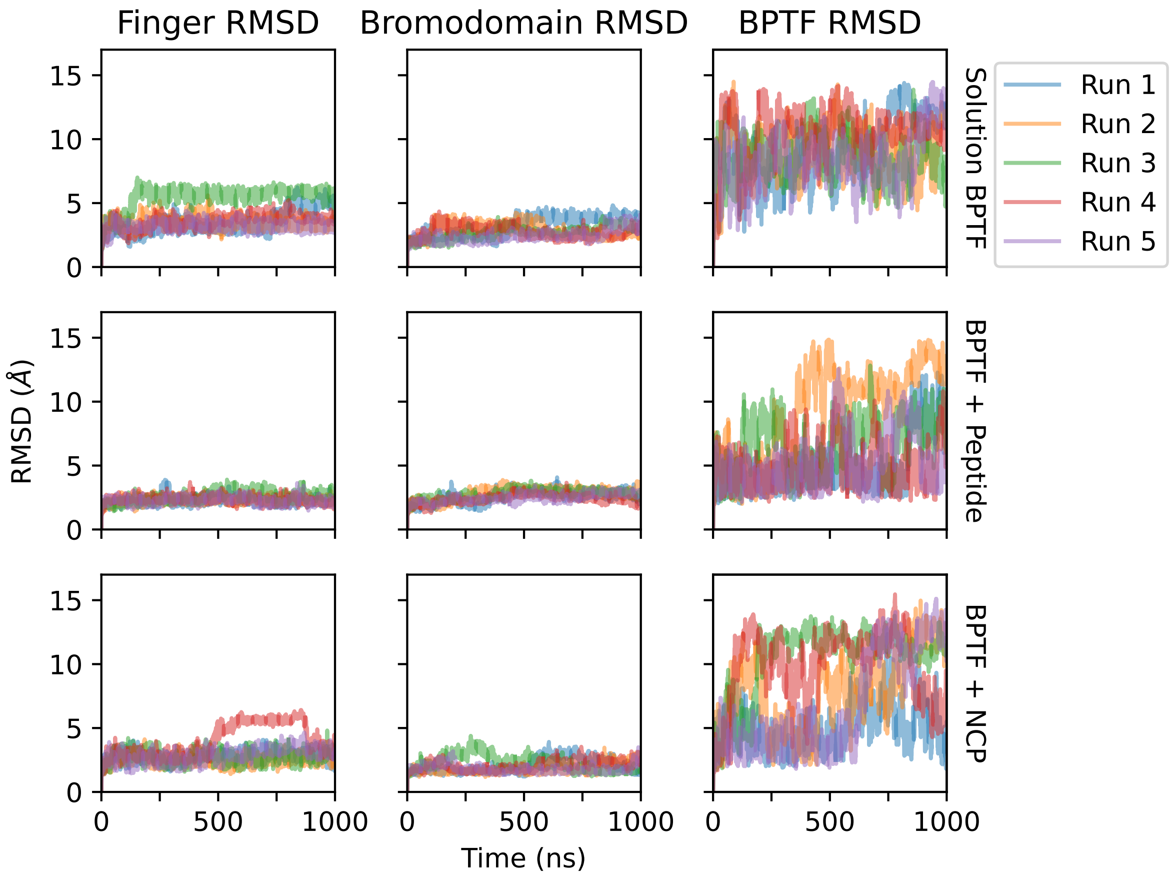


Figure S17: Root mean squared deviation by region of BPTF from the first frame. While the PHD finger (here taken as residues 1 through 59) and bromodomain (residues 66 through 168) are relatively stable, the loop region between them enables a hinge-like motion, leading to high mobility of each domain relative to the other.

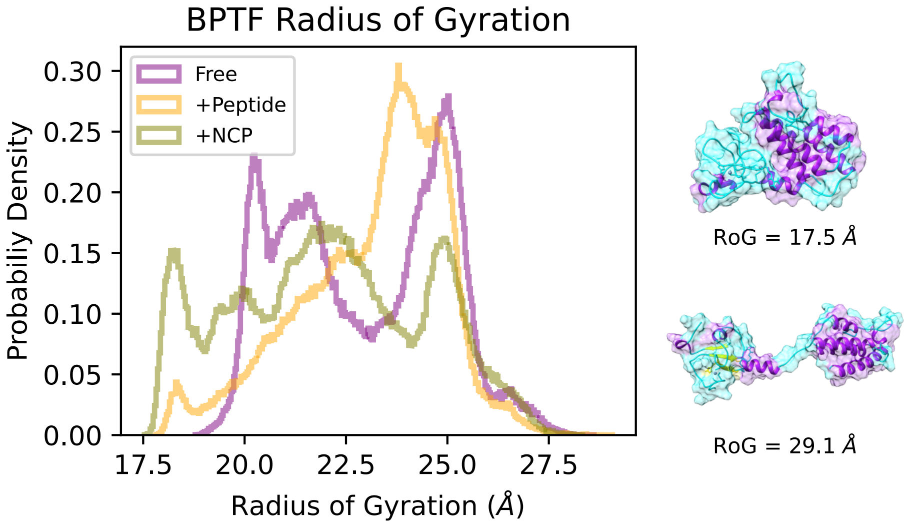


Figure S18: Left: The radius of gyration of BPTF as measured across free, peptide bound, and NCP bound systems. While all contexts see large variation, short peptide binding rigidifies the protein in a way not represented by free and NCP-bound BPTF. Right: Sample conformations of BPTF in varying levels of compactness with the corresponding radius of gyration.

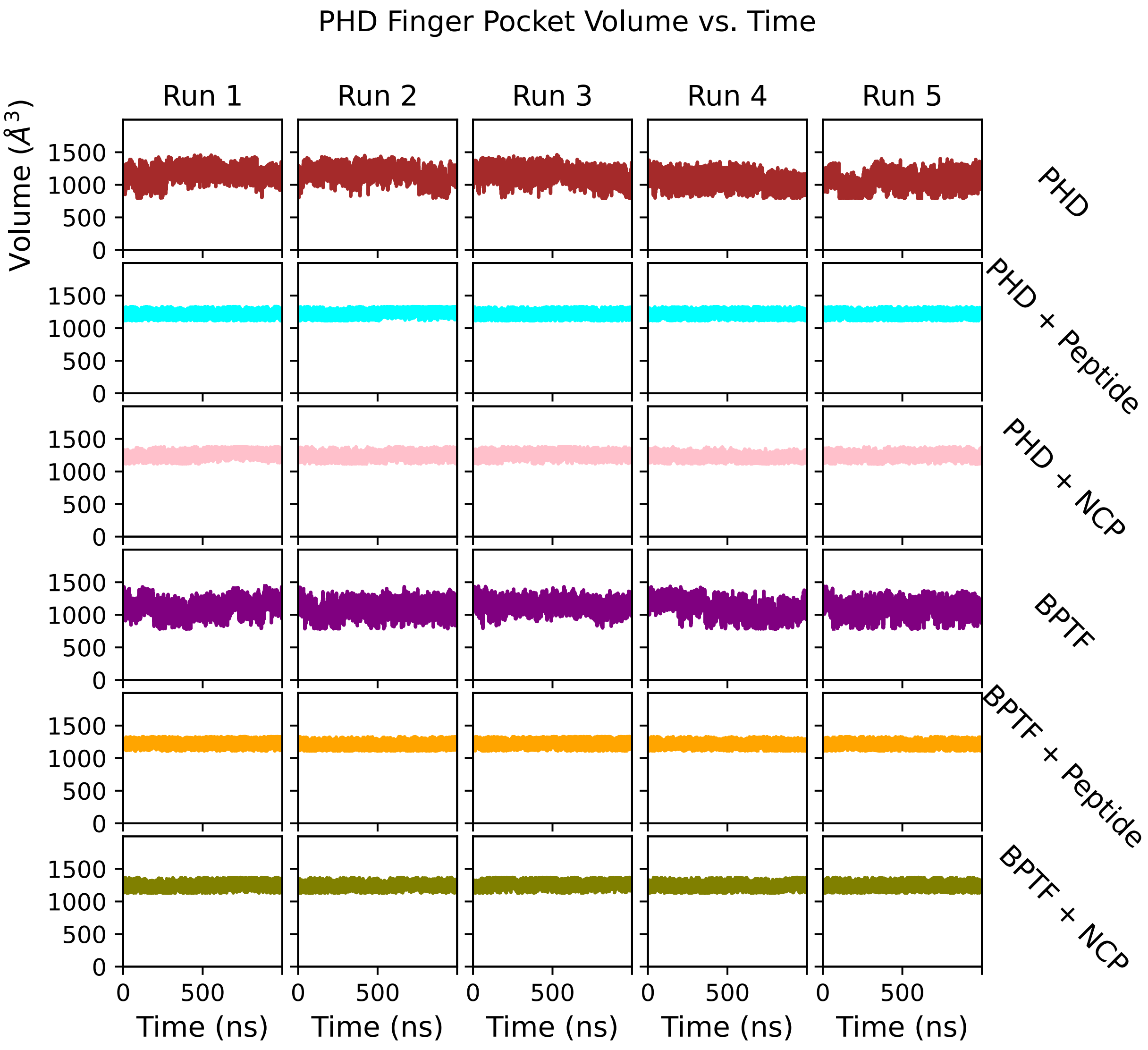


Figure S19: Time series of the PHD finger binding pocket volumes of all systems. Opening and closing of the pocket represents a characteristic motion of the PHD finger, and H3 binding serves to stabilize this motion and hold the pocket in a narrower, more open range over time.
